# Spatial remapping in the subicular complex and entorhinal cortex follows low-dimensional geometric principles

**DOI:** 10.64898/2026.05.10.724068

**Authors:** Shai Abramson, Daniel Zur, Gaya Tzadok, Shachaf Kolan, Shiladitya Laskar, Ohad Rechnitz, Vijay Balasubramanian, Genela Morris, Hadas Benisty, Dori Derdikman

## Abstract

Neurons in the hippocampal formation are part of the brain’s cognitive map, representing the spatial structure of the environment through coordinated activity across place cells, grid cells, border cells, head-direction cells and others^1–5^. Although remapping between environments has been extensively documented^6–9^, it remains unknown whether transitions between maps reflect unconstrained reorganization or obey a systematic transformation principle. To address this question, we recorded large neuronal populations from the subicular complex and entorhinal cortex in awake, behaving mice navigating environments spanning a wide range of geometries. We asked whether spatial representations across rooms could be related through a shared class of coordinate transformations. Despite pronounced heterogeneity and apparent randomness in single-cell remapping, population-level decoding across environments demonstrated a consistent low-dimensional affine transformation of coordinates, comprising rotation, scaling, shear, reflection, and translation. Thus, what appears as complex remapping at the level of individual neurons reduces to a compact geometric rule at the level of neural assemblies. These results indicate that the hippocampal formation maintains a structured internal coordinate template that is flexibly tailored to environmental geometry. This may serve as the organism’s internal model of space.

## 1 Introduction

The hippocampal formation has been known for years to play a pivotal role in the brain’s cognitive map in general, and specifically in constructing a representation of space^10–13^. A major unanswered question is how the system generalizes between different environments. When moving between domains, the neural representation changes, a phenomenon known as *remapping* ^6,9,14^. However, the rules governing remapping of the population are unknown. We asked whether we could find some form of representational alignment of the neural code between different environments. To determine whether such an alignment existed, we recorded hundreds of neurons simultaneously in multiple rooms of various shapes while mice performed a guided exploration task. We recorded mostly from the medial entorhinal cortex (MEC) and the subicular complex (SUBC), which contain cell types that serve as core building blocks of the cognitive map in the hippocampal formation^2–4,15–19^. We decoded spatial location from population activity across different environments and applied our analysis to determine the transformation of spatial map coordinates between rooms. We found that coordinate transformations during remapping were often well described by low-dimensional affine transforms: combinations of rotation, reflection, scaling, shear, and translation, revealing a principled geometric organization at the assembly level. Thus we were able to reveal a global principle of representation alignment^20,21^ between different environments in the cognitive map.

## 2 Results

To probe population-level remapping across distinct spatial contexts, we recorded large-scale neuronal activity from freely moving mice with Neuropixels probes targeted to medial entorhinal cortex (MEC), subiculum (SUB) and presubiculum (PRE) (Fig. 1a–d, Supplementary Fig. **??**a– d). Mice explored a custom-built two-room arena (Fig. 1e) during daily sessions consisting of three consecutive 20-min epochs: exploration of room A, transition to room B, and a return to room A (A–B–A^′^; Fig. 1f). Across recording sessions, we systematically varied room geometries, generating varied pairs of environments with distinct combinations of straight and curved boundaries (Fig. 1g). Overall, We recorded 71 sessions from 6 animals, yielding a total of 9,009 single units in all sessions, out of which 3,855 units were in MEC, 3,070 units were in SUB and 965 units were in PRE. To augment the number of cells for population analyses we grouped the SUB and PRE together into the SUBC region (Table **??** and Methods, Section 4.4).

**Figure 1:**
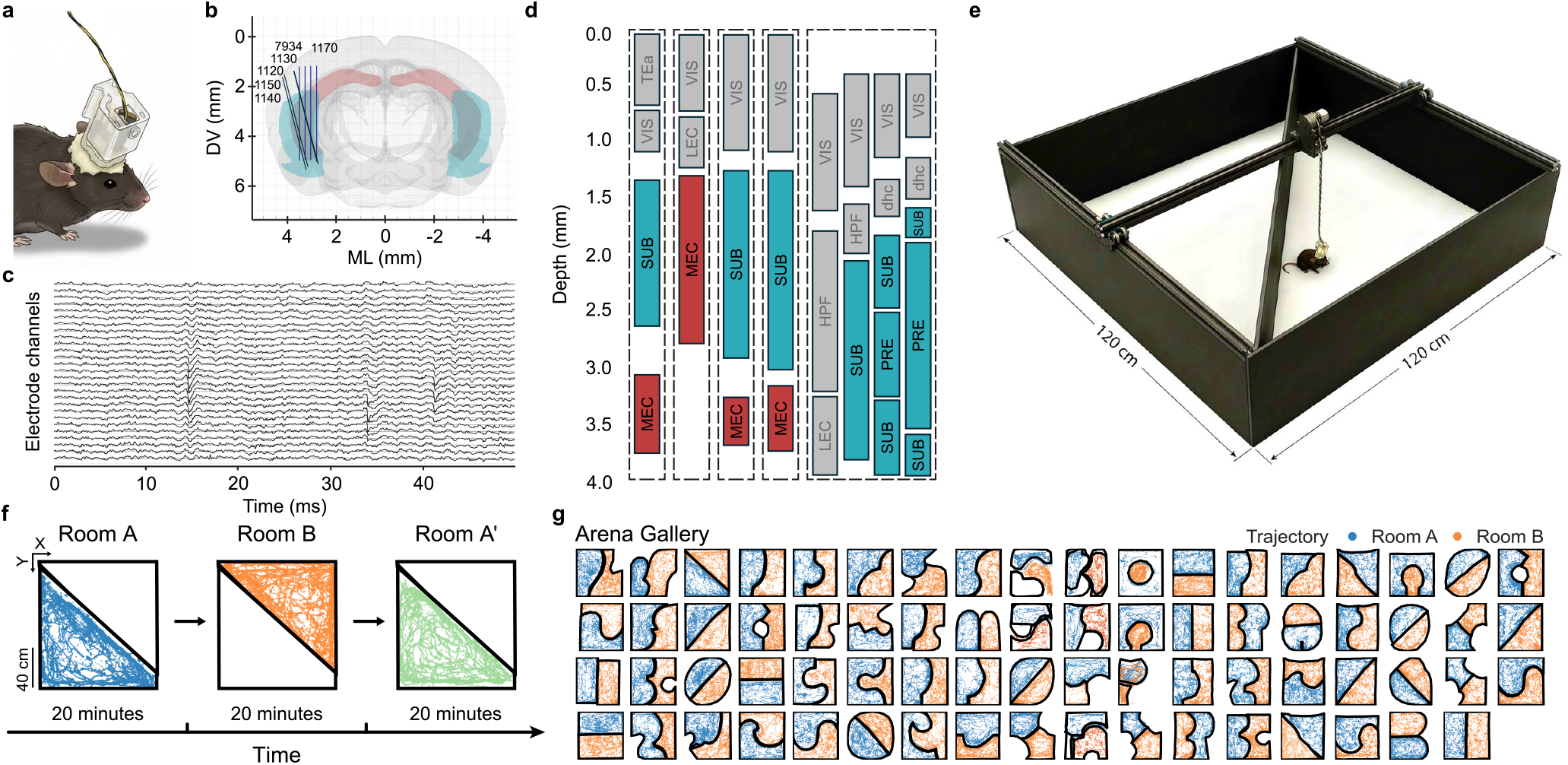
Large-scale recordings in the hippocampal formation during spatial navigation across distinct room geometries. **(a)** Schematic of a mouse implanted with a Neuropixels probe for chronic extracellular recording during free exploration. **(b)** Coronal brain section showing reconstructed probe trajectories. Neuropixels 2.0 multi-shank probes are shown in blue and Neuropixels 1.0 single-shank probes in black; animal numbers are indicated next to each probe. (c) Representative extracellular voltage traces from adjacent channels showing simultaneous spiking activity (scale bars, 10 ms and 100 µV). **(d)** Distribution of recording sites along the probe shanks spanning cortex, subiculum (SUB, cyan), presubiculum (PRE, cyan), and medial entorhinal cortex (MEC, red); Regions not included in the analysis due to small numbers of cells are marked in gray. From left to right, the schematics correspond to mice 1140 and 1150 (same coordinates), 1120, 7394, 1130, and 1170 (four shanks). **(e)** Custom 120 × 120 cm two-room arena with a motorized tether system for unrestricted exploration. **(f)** Behavioral protocol: mice explored room A (0–20 min), then room B (20–40 min), and then returned to room A^′^ (40–60 min). Traces show representative trajectories in room A (blue), room B (orange), and room A′ (green). **(g)** Room geometries comprising diverse combinations of straight and curved boundaries in symmetric and asymmetric layouts.

### Diverse functional spatial cell types

We first classified neurons by their tuning properties in each room to ask whether functional cell identity was preserved across environments. Units were classified according to quantitative tuning metrics, with significance assessed against shuffle-based null distributions (see Methods, Section 4.4.2). In both MEC and SUBC, we identified different functional classes, including allocentric border cells (ABCs; Fig. 2a), egocentric border cells (EBCs; Fig. 2b), head-direction (HD) cells (Fig. 2c), spatially selective cells with high spatial information (Fig. 2d), and speed-modulated cells (Fig. 2e). Many neurons exhibited conjunctive tuning, belonging to more than one functional class.

**Figure 2:**
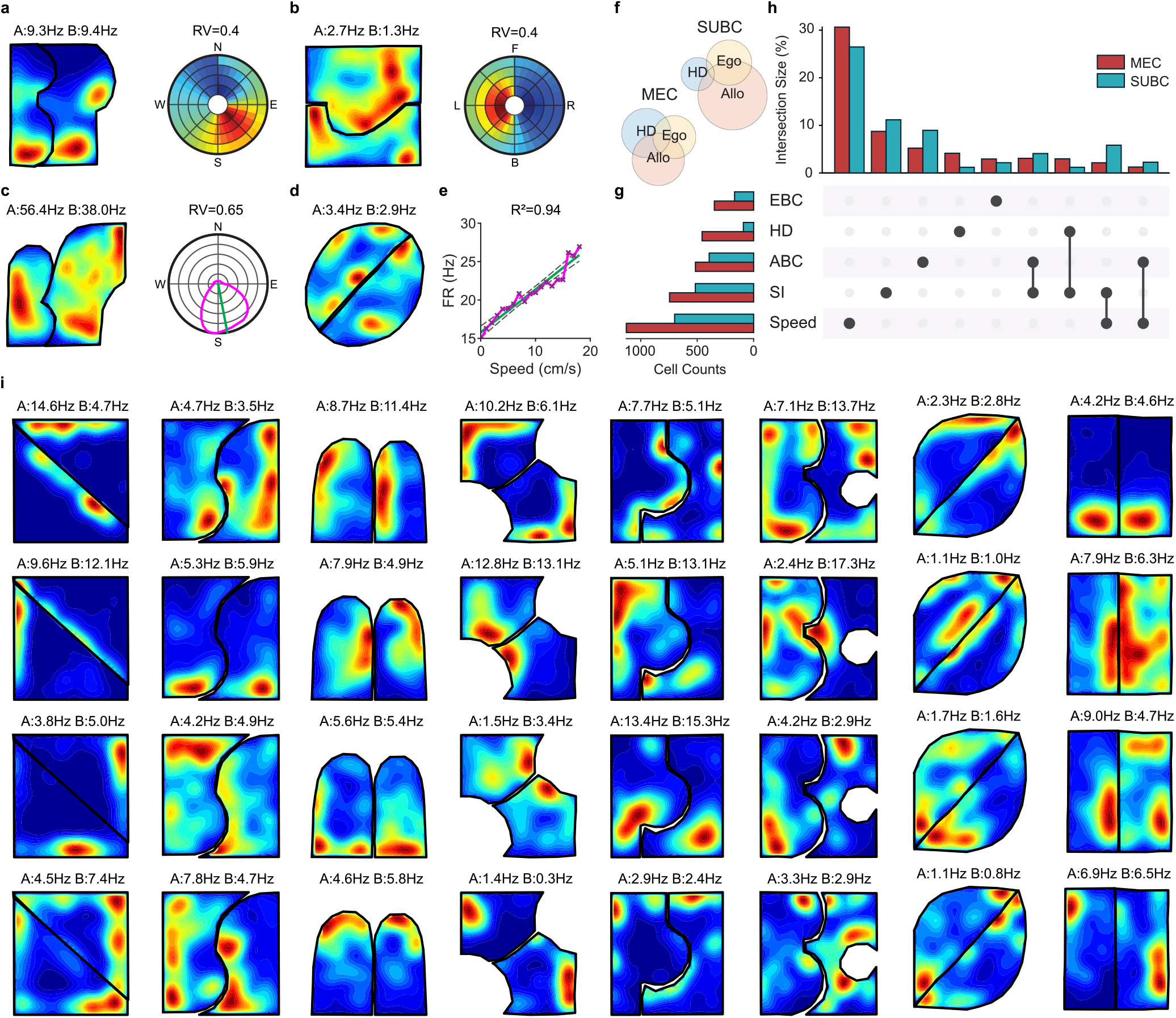
Diversity of spatial cell types in the SUBC and MEC. **(a)–(e)** Representative examples of functional cell types. Each panel shows a spatial rate map (left) and a relevant tuning curve (right). **(a)** Allocentric border cell (ABC) with boundary-aligned firing fields (Rayleigh vector (RV) ≥ 0.3). **(b)** Egocentric border cell (EBC) tuned to boundary distance and angle in a head-centered frame (RV ≥ 0.3). **(c)** Head-direction (HD) cell with allocentric preferred direction (RV ≥ 0.3). **(d)** Spatially modulated cell with high spatial information (SI *>* 1). **(e)** Speed cell with linear relationship between firing rate and running speed (*R*^2^ *>* 0.6). **(f)–(h)** Population summaries and overlap among functional cell types in the SUBC and MEC. Classification required metric significance (*P <* 0.05) relative to a shuffled null distribution. **(f)** Venn diagrams showing population overlap of ABC, EBC, and HD properties in SUBC and MEC. **(g)** Proportions of cells classified into each functional category by region. **(h)** An UpSet plot quantifying conjunctive coding (multi-modal tuning) across the population. **(i)** Example Allocentric Border Cells (ABC) rate maps for rooms A and B across sessions with differing room geometries, illustrating persistent boundary anchoring. Peak firing rates (Hz) are indicated for each room.

Overlap among cells classified as one of the directionally-oriented classes (ABCs, EBCs, HD cells) revealed substantial conjunctive coding in both MEC and SUBC regions (Fig. 2f–h). Across the broader population, speed-modulated cells were most prevalent in both MEC and SUBC, followed by ABCs, EBCs, and HD cells. Specifically, we identified *n* = 158 exclusively significant ABCs in MEC and 151 in SUBC; 85 EBCs in MEC and 38 in SUBC; and 253 high– spatial-information cells in MEC and 197 in SUBC. These cell classes enabled comparison of firing-rate maps across room geometries (Fig. 2i).

We next examined the stability of representations across rooms. Comparing significant units between epochs (A vs. B and A vs. A^′^) indicated that most cells retained their classification (see Methods Section 4.4.3 and Supplementary Fig. **??**). Classification stability varied by cell class, exceeding 87% for high-spatial-information cells and 98% for speed cells, and remaining above 53% of ABCs, EBCs, and HD cells.

For neurons that remained significantly directional in both rooms (ABCs, EBCs, HD cells), we quantified changes in preferred orientation (see Methods, Section 4.4.4). The resulting distributions of the angle changes were broad (Supplementary Fig. **??**). ABCs showed absolute angle differences spanning the full 0–180° range (*n* = 146 and 209 cells for A–B and A–A^′^, respectively) and EBCs showed a similarly wide spread (*n* = 44 and 57 for A–B and A–A′). HD cells were too few in our recordings (*n* = 13 and 14 for A–B and A–A′) to reliably characterize their angular distribution. Thus, functional class identity was often preserved across rooms, whereas the preferred orientation among directional cells frequently shifted.

### Neural network decoder uncovers continuous geometric transformations between environments

To investigate population remapping without imposing linear or time-invariant assumptions, we developed an unbiased analysis framework that captures nonlinear and temporally structured relationships across contexts. The pipeline comprised two components: a recurrent temporal encoder that maps population activity across all rooms to a shared latent space, and a room-specific decoder that transforms the latent coordinates to physical positions within each room (see Methods, Section 4.5, Fig. 3a). Note that this is reminiscent of previous models of the hippocampus as a recurrent encoder of sensory and other inputs with emergent place and time cells^22–25^.

**Figure 3:**
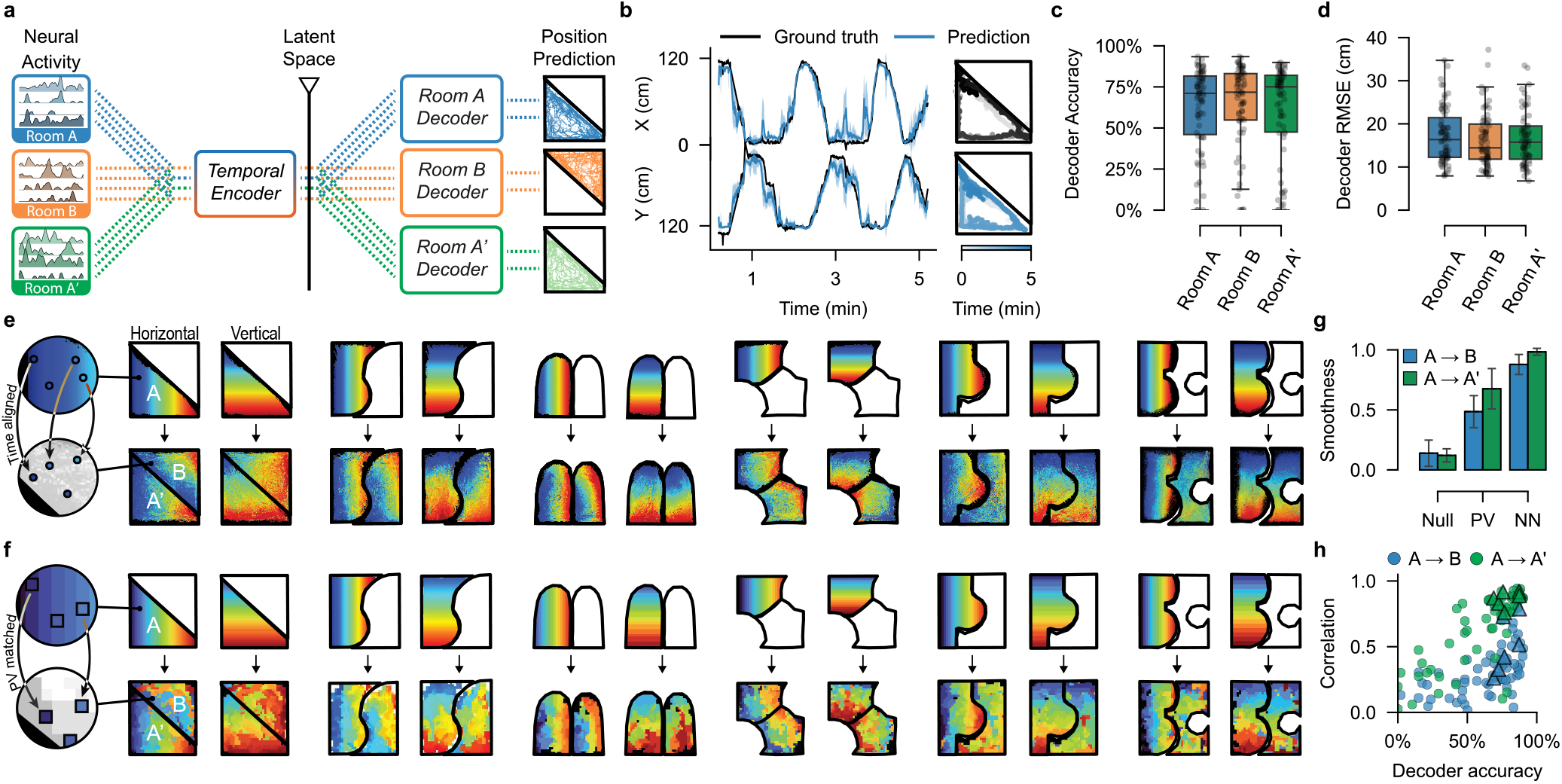
Continuous remapping of spatial representations revealed by a neural network position decoder. **(a)** Model architecture. A shared temporal encoder maps population activity sequences from all rooms (A, B, A′) to a common low-dimensional latent space. Distinct decoder heads map this latent state to position in each room’s coordinate frame. The encoder was trained on activity traces from rooms A and B, while each decoder head was trained solely using data from its respective room (blue: A; orange: B; green: A′). **(b)** Example: ground truth coordinates (black) and the decoder predictions (blue; mean ± standard deviation across repetitions) during 5 minutes. Right: Spatial 2D view of the trajectory color-coded by time. **(c)** Decoding accuracy 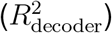 across sessions (*N* = 71) for Rooms A, B, and A′. Boxplots show median and IQR; dots represent individual sessions. Median accuracy exceeds 0.7 across contexts. (d) Position decoding error (RMSE, cm) for the same sessions. Median error typically ranges from 10 to 20 cm. **(e)** Visualization of decoder-based remapping. Schematic (left): time-aligned correspondence between decoded positions. Main gallery: Source-room positions colored by horizontal (*x*) or vertical (*y*) coordinate (top), mapped to decoded target-room locations (bottom), revealing smooth, geometry-preserving gradients. **(f)** Comparison with population vector (PV) remapping. Schematic (left): PV-based spatial bin correspondence. Main gallery: Equivalent gradient analysis using PV correlations. Mappings are noisier but exhibit similar geometric structure. **(g)** Smoothness quantification. Comparison of spatial mapping smoothness (Laplacian-based score, see Methods, Section 4.6.3) across three conditions: Null (random bin mapping), PV (population-vector mapping), and NN (decoder-based mapping), for A →B (blue) and A →A′ (green). **(h)** Geometric agreement between NN and PV mappings quantified by Pearson correlation between target coordinates inferred by the NN decoder and PV analysis, plotted versus mean decoder accuracy (average 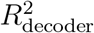 across rooms) for mappings A→B (blue) and A→A′ (green). Triangular markers indicate the example sessions displayed in (e) and (f).

We designed this architecture so it would: 1) provide a data-driven mapping from population dynamics to position; 2) disentangle the shared temporal code from room-specific spatial information; 3) allow us to explore how population activity related to a particular room would be decoded in any of the three rooms (A, B and A’). First, we evaluated the performance of our pipeline in decoding the animal’s position from population dynamics, on a per-room basis (models were trained using 10-fold cross-validation with 10 independent repetitions, see Methods, Section 4.5, and Supplementary Fig. **??**c). Example trajectories demonstrate reliable tracking of position over time and accurate reconstruction of spatial paths (Fig. 3b). Across all sessions (*N* = 71), the model achieved high decoding accuracy (median 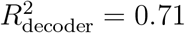, with inter-quartile range [IQR] [0.46–0.82] for room *A*, 0.72 [0.55–0.83] for room *B*, and 0.75 [0.48–0.82] for room *A*^′^; Fig. 3c), with correspondingly low position decoding error (16.4 cm [12.2–21.4] for room *A*, 14.4 cm [11.8–19.9] for room *B*, and 15.7 cm [11.8–19.5] for room *A*^′^; Fig. 3d), confirming that the model reliably maps population activity to spatial position. Note that this architecture outperformed both linear and non-linear time-independent baselines (Supplementary Fig. **??**h).

### The shared latent space is low-dimensional and room-invariant

To determine whether the shared temporal encoder produces a compressed representation for neural dynamics without introducing contextual separation, we quantified the variability of raw neural activity traces relative to the latent representation using Principal Component Analysis (PCA). For the raw data, 71 ± 37 principal components were required to capture 95% of its variance, while the latent representation required a much smaller number of 9 ± 2 components to capture the variability of the 64-dimensional latent representation (mean ± s.d. across sessions; Supplementary Fig. **??**e). Crucially, a linear Support Vector Machine (SVM) classifier based on the first three principal components of this compressed latent representation was unable to discriminate environmental context better than when based on the raw activity (Supplementary Fig. **??**e). Overall, these results suggest that the shared latent space effectively reduced the dimension of the representation while maintaining room-invariance.

### Neural activity maps continuously between different spatial contexts

We next used our architecture to determine how the encoding of neural dynamics related to one environment is transformed to represent another. We passed the neural activity recorded in a source room through the temporal encoder, thereby generating latent trajectories in the shared space. These latent coordinates were then decoded using both the source-room and target-room decoder heads. In this way, the same underlying neural activity produced two temporally matched spatial sequences: the estimated position of the mouse in the source room and the corresponding position decoded in the target room’s frame of reference. This approach allowed us to define, at each time point, a pair of corresponding spatial coordinates that link the two environments (see Methods, Section 4.6.1; Supplementary Fig. **??**i).

We visualized the spatial structure of this remapping by coloring the source-room decoded positions by their horizontal (*x*) or vertical (*y*) coordinates, and projecting these colors onto the corresponding decoded positions in the target room (Fig. 3e). The obtained mappings showed that spatial representations undergo structured geometric transformations. Although the analysis was highly nonlinear and data-driven, the resulting geometric transformations were usually simple, as quantified in the following sections.

### Decoder-derived transformations match population vector analysis

To confirm that the visualized geometric structure is invariant of the decoding method, we next asked whether the geometric structure captured by the neural network (NN) decoder could also be detected by a conventional population-vector (PV) correlation analysis. For each spatial bin in one room, we identified the bin in the other room that maximized the Pearson correlation between their PVs, and averaged these maxima across bins to yield a session-level summary statistic (see Methods, Section 4.6.1). Observed maximum PV correlations strongly exceeded the shuffle-based null correlations in 92.3% of sessions for each room combination (A–A′, A–B, and B–A′), (median *P <* 0.001; Methods, Section 4.6.1; Supplementary Fig. **??**a,b).

To systematically compare this approach to our NN mapping, we constructed PV-based correspondences between rooms by matching spatial bins via maximum PV correlation (Fig. 3f; see Methods, Section 4.6.1). We then discretized the continuous NN decoder output to match the PV spatial binning for direct comparison (see Methods, Section 4.6.3). We quantified the preservation of local spatial structure by computing a graph-based smoothness score (Eq. 2; see Methods, Section 4.6.3). This approach evaluates whether neighboring bins in the source room map to contiguous locations in the target room by measuring the local coordinate variation on a graph. We evaluated the smoothness score for each mapping method (NN and PV) across distinct rooms (A → B) and identical room geometries (A → A′ as a positive control), comparing the results to a null distribution of random bin assignments. We found that NN mappings were the smoothest (mean smoothness score ±s.d.: 0.98 ± 0.03 for A → A′, 0.88 ± 0.08 for A → B), followed by PV (0.68 ± 0.17 for A →A′, 0.49 ± 0.13 for A →B), with the Null baseline significantly lower (0.12 ± 0.06 for A → A′, 0.14 ± 0.11 for A → B; Fig. 3g). Both PV and NN mappings were significantly smoother than the null (*P <* 0.001), confirming that the observed transformations preserve local neighborhood relationships. Furthermore, NN mappings were significantly smoother than PV mappings (*P <* 0.001), reflecting the NN’s ability to recover a more coherent, continuous map. The maps obtained by the NN and PV methods were well correlated, and the correlation increased with decoding accuracy (Fig. 3h). We conclude that the coordinate transformations obtained by the NN and PV methods were similar, suggesting that it was independent of the method of derivation.

### Coordinate transformation possesses affine properties

Visual observation of the obtained coordinate mappings (Fig. 3e) revealed a large proportion of transformations that could be described by a rotation, stretch, and reflection of the coordinates. We therefore asked to what extent the transformation in both methods could be described as affine, namely comprising translation, rotation, possible reflection, scaling, and shear (Fig. 4c).

**Figure 4:**
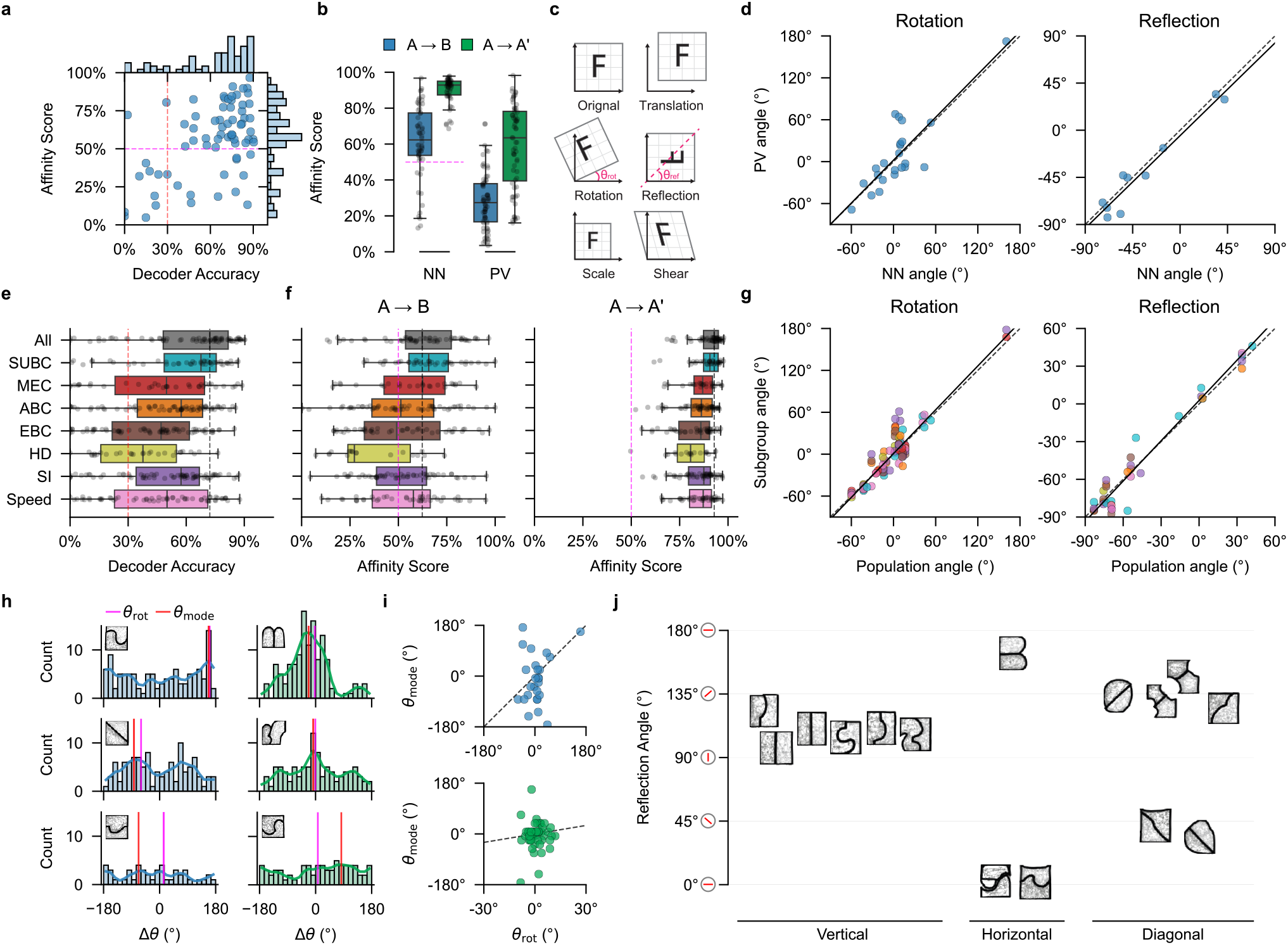
Spatial remapping follows low-dimensional affine transformations anchored to environmental geometry. **(a)** Joint distribution of decoder accuracy and affinity score for A→B mapping. Marginal histograms show distributions. Dashed lines: inclusion thresholds 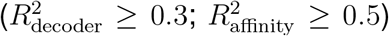. **(b)** Comparison of affinity scores between neural-network (NN) and population-vector (PV) methods for A→ B and control A →A′ mappings. Boxplots show the median and IQR, with individual session scores overlaid. **(c)** Schematic of 2D affine transformation components. **(d)** Agreement of transformation angles between decoder (NN) and population-vector (PV) methods. Dashed lines: identity; solid lines indicate circular-linear regressions (*P <* 0.001) for rotation angles (left; slope 1.02, RMSE 26.8°, *N* = 23) and reflection angles (right; slope 0.98, RMSE 10.0°, *N* = 10). **(e)** Decoder accuracy among neural subpopulations: full population (‘All’), regions (MEC, SUBC), and functional types (ABC, EBC, HD, SI, Speed). Dashed lines mark the median for the full population as baseline (black) and the decoder accuracy threshold (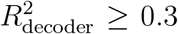 red). **(f)** Affinity scores across neural subpopulations (A→B and A→A′). Dashed lines mark the full-population median (black) and the affinity score threshold (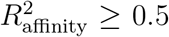; red). **(g)** Consistency of transformation angles across subpopulations. Dashed lines: identity; solid lines indicate significant circular-linear regressions (*P <* 0.001) for rotation angles (left; slope 0.98, RMSE 23.7°, *N* = 131) and reflection angles (right; slope 1.02, RMSE 5.7°, *N* = 54). **(h)** Per-session distributions of ABC angular shifts (Δ*θ*_*i*_) in rotation sessions (left: *A* →*B*; right: *A*→ *A*^′^). Vertical lines indicate the global rotation angle (*θ*_rot_, magenta) and the circular peak mode of single-cell shifts (*θ*_mode_, red). **(i)** Session-level conformity. Scatter plots relating the single-cell shift mode (*θ*_mode_) to the global rotation angle (*θ*_rot_) for *A* →*B* (top; *N* = 26 sessions) and *A* →*A*^′^ (bottom; *N* = 49 sessions). Dashed lines: identity. **(j)** Alignment of reflection axes with room geometry. Reflection axes are grouped by the orientation of the shared wall (vertical, horizontal, or diagonal (±45°)). Vertical lines indicate the expected axis direction (red).

To quantify this, we fitted a global affine transformation to the cross-room position correspondences. We defined an affinity score as the coefficient of determination (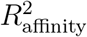, ranging between 0 and 100%) of this fit (see Methods, Section 4.6.4). This score evaluates how well the complex spatial remapping is captured by simple geometric operations, with higher values indicating that the transformation between environments is predominantly affine.

The joint distribution of decoding accuracy and affinity scores (Fig. 4a) effectively partitioned the data into distinct regimes. First, in the low-accuracy regime, characterized by low decoding accuracy 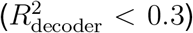, the affinity scores were predominantly low, reflecting the inherent impossibility of recovering precise geometric transformations from noisy position estimates. Second, once decoding became reliable 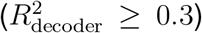, the distribution bifurcated: 58 sessions (83%) reached high accuracy, of which a minority remained in the low-affinity quadrant, while the majority (81%; *N* = 47) converged into a distinct high-affinity cluster 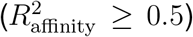. This separation indicates that whenever the spatial code is robust, the remapping between rooms is predominantly affine. In subsequent analyses in this section, we quantified the affinity of these transformations for all reliably decoded sessions (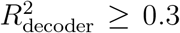; *N* = 58 sessions for *A* → *B, N* = 54 sessions for *A* →*A*^′^ and *B* →*A*^′^). To characterize the specific parameters of the transformations (such as rotation and reflection angles), we restricted our subsequent analyses to the high-confidence regime (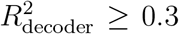 and 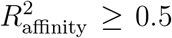) where the geometric structure is well-defined and can be reliably estimated.

Affinity scores for mapping between rooms (A →B mappings) were significantly higher for NN than those derived from PV-based correspondences (0.62 [0.54–0.77] for NN vs. 0.27 [0.17– 0.38] for PV; Fig. 4b), reflecting the NN decoder’s superior noise robustness. Similarly, for returning to the same room (A→ A′), the affinity scores were higher for the NN (0.93 [0.87–0.95] for NN vs. 0.64 [0.40–0.78] for PV). Despite these differences in fit quality, both methods showed the same qualitative pattern: A →A′ mappings yielded higher affinity than A →B mappings.

We then compared the affine parameters recovered by the two approaches. For both NN and PV-based correspondences, we decomposed the affine transformations into rotation, reflection, scaling, shear, and translation components (Fig. 4c). For this comparison, we included only sessions in which both methods agreed on whether the affine transformation was a rotation or a reflection (70% agreement: *N* = 23 rotation, *N* = 10 reflection; see Methods, Section 4.7). We found that rotation angles (*θ*_rot_) were strongly correlated between methods (circular-linear regression: *R*^2^ = 0.61, *P <* 0.001; Fig. 4d left), indicating that both approaches recover similar remapping rotations. For sessions with reflection components, reflection-axis angles (*θ*_ref_) showed good alignment between methods (*R*^2^ = 0.90, *P <* 0.001; Fig. 4d right). This cross-method agreement confirms that the geometric transformations resolved by the NN decoder reflect fundamental population-level remapping that remains detectable, despite increased noise, using more traditional PV-based analysis.

### A unified geometric transformation governs remapping across diverse neural subpopulations

We next asked whether the geometric structure of remapping discussed above reflects a global network property or is specific to particular functional modules. We retrained our room decoding architecture (Fig. 3a) separately for each subpopulation defined by brain region (SUBC, MEC) and cell type (ABC, EBC, HD, SI, Speed, see Methods, Section 4.8). SUBC cells enabled decoding accuracy similar to that of the full population, albeit with fewer cells (0.68 [0.49–0.76] for SUBC vs. 0.72 [0.48–0.82] for the full population, Fig. 4e). The MEC cells, however, yielded somewhat lower values (0.50 [0.23–0.70], Fig. 4e). Across functional cell types, ABC and SI cells yielded the best decoding (0.57 [0.35–0.69] for ABC, 0.57 [0.34–0.67] for SI), followed by Speed (0.50 [0.23–0.71]), EBC (0.47 [0.22–0.62]), and HD cells (0.38 [0.16–0.55]).

Moreover, affinity scores for *A* →*B* remained stable across subpopulations (median 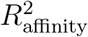 was 0.62 [0.54–0.77] for the full population, 0.66 [0.55–0.76] for SUBC and 0.63 [0.43–0.74] for MEC, and 0.49–0.58 across ABC, EBC, SI, and Speed, with HD lower at 0.27 [0.24–0.56], Fig. 4f). Control *A* → *A*^′^ mappings yielded high affinity across all groups (medians 0.81–0.93). Most sessions exceeded the affinity threshold 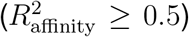 for the full population (81%), for SUBC (81%), and for MEC (68%), with moderate rates for specific cell types (50–59%), indicating that the geometric structure of cross-room transformations is a robust property shared throughout diverse cell types and brain regions. The affine parameters learned per subpopulation were consistent with those of the full population, both for rotation (*R*^2^ = 0.68, *P <* 0.001) and reflection (*R*^2^ = 0.97, *P <* 0.001; Fig. 4g).

Finally, we tested whether single-cell tuning shifts align with the population-level transformation. To this end, we compared each cell’s ABC directional shift with the session-level rotation angle (*θ*_rot_), restricting the analysis to sessions best described by a rotation (*N* = 26 for *A* →*B* and *N* = 49 for *A* →*A*^′^; see examples in Fig. 4h). Defining alignment as an angular difference of ≤30° between the predominant single-cell shift angle (the circular mode of individual shifts, *θ*_mode_) and the global affine transform (*θ*_rot_), we found that 9 of 26 *A* →*B* sessions (34.6%) and 32 of 49 *A* →*A*^′^ sessions (65.3%) showed alignment. Generally, we conclude that the transformation of single cells was heterogeneous in comparison to the global transformation in the *A → B* case, and less so when returning to the same room (Fig. 4i).

### Spatial reflections are anchored to environmental geometry

In a subset of sessions, the best-fitting affine transformation included a reflection (*N* = 15 sessions; see Methods, Section 4.11). To test whether reflections are geometrically constrained, we compared the orientations of reflection axes to the direction of the wall shared between the rooms. We defined alignment as an angular difference ≤30° relative to the shared wall (see Methods, Section 4.11). Reflection axes aligned systematically with room geometry in 14 out of 15 sessions (93.3%; Fig. 4j). Horizontal and diagonal geometry sessions deviated by at most 15.9° from the expected axis (*N* = 9), while vertical geometry sessions (*N* = 6) ranged up to 33.5°, with 5 of 6 falling within 21.1°. This systematic alignment indicates that reflections are anchored to environmental boundaries rather than constituting arbitrary transformations.

## 3 Discussion

Spatial representations in the hippocampal formation change substantially across environments^6^ and tasks^26^, yet navigation remains coherent. We recorded large populations in the MEC and subicular complex during exploration of systematically varied room geometries. We chose to start with those regions because there is evidence that they show less change than CA1 during remapping situations^8,15,19,27–31^. We used this activity to decode position within these rooms. A consistent principle emerged: the cross-room coordinate transformation was well captured by a low-dimensional affine mapping (rotation, reflection, scaling, shear, and translation). This suggests that spatial remapping in the MEC and SUBC reflects a structured coordinate transformation rather than an arbitrary reassignment of spatial relationships. This is in contrast to remapping in the hippocampus proper, which appears to be random at the individual cell level^7,32^, but is consistent with accounts pointing to a low-dimensional manifold during the transformation in these regions^9,20,33–36^. Methodologically, to extract these continuous transformations we relied on our custom motorized tethering system, which mitigated the behavioral sampling biases typical of free exploration.

Two complementary analyses, a decoder approach and a population-vector approach^7,20,32,37^, recovered similar affine parameters. The methodological convergence confirms that the geometric structure that we have revealed is a property of the underlying neural manifold, rather than an analytical artifact. Despite the decoder’s flexibility to learn arbitrary nonlinear mappings, simple affine transforms were sufficient to capture remapping between environments. These findings suggest that remapping follows low-dimensional geometric principles rather than random reorganization. Our results may be related to foundational observations of structured grid cell phase shifts and modularity, which theoretically permit global affine transformations of the spatial map^8,38,39^. Furthermore, previous work has demonstrated that the correlation structure of these grid networks remains robustly preserved even when external spatial inputs are disrupted^40–43^.

While this may be the case, affine transformations have not yet been shown systematically in the case of grid cells on multiple, diverse-shaped rooms. Rather, there are known cases where grid manifolds deform from affinity when room shapes change drastically^38,44,45^, possibly because of the effects of interactions with border cells^46,47^. Our results are furthermore in dialogue with recent manifold analyses demonstrating that hippocampal subpopulations undergo independent transformations^48^ and conserve population geometry despite pronounced single-cell remapping^49^.

By presenting these affine transformations in the MEC and subicular complex, our results demonstrate how boundary-anchored codes^4,19,50^ can undergo structural transformations between differently shaped enclosures. This population-level transformation was broadly expressed across anatomically and functionally defined subpopulations. At the single-cell level, however, neurons displayed highly heterogeneous shifts. This interplay between single-cell variability and ensemble stability mirrors a well-known property of representational drift^51–53^: individual cells may reorganize to multiplex contextual or sensory variables, while the population retains a coherent spatial map.

A particularly informative subset of sessions exhibited reflection components that aligned with the wall orientation shared between the two rooms. This suggests that the shared boundary serves as a “geometric hinge”, anchoring the transformation between rooms and points to the existence of a brain geometric module^54^, and that local geometry dictates the structural limits and alignment of spatial codes^30,55–57^. Moreover, this aligns with findings that the broader hippocampal-parahippocampal circuit is inherently organized to compute continuous coordinate transformations relative to spatial boundaries^58^. Upon exposure to a new room^6,59^, rather than constructing an orthogonal map de novo, the network appears to actively align its internal coordinate frame with the new space via template matching. This is consistent with evidence that the hippocampus maintains preconfigured network states that are rapidly recruited and selected to represent novel environments^60,61^.

Our findings provide a compact geometric vocabulary for spatial representation. Rather than treating remapping as a categorical switch, we show that cross-environmental changes are governed by a well-defined family of geometric operations acting on a population code. Our work provides a population-level framework for analyzing coordinate transformations in the brain. The advantage of having such a canonical geometric transformation between rooms at the population level is that local relations structure, and hence computations that depend on such relations structure, can be left unchanged. Beyond spatial navigation, these results suggest a general mechanism by which neural systems may preserve relational structure while adapting to new contexts, with implications for theories of neural computation and representation alignment^20,21^.

## 4 Methods

### 4.1 Animal Preparation and Surgery

#### 4.1.1 Mice

Electrophysiological recordings were performed in six adult male C57BL/6J wild-type mice, aged 8–20 weeks at the time of surgery. Mice were housed individually in transparent Perspex cages with sawdust bedding under controlled ambient conditions (20–24 °C), with ad libitum access to food and water on a 12-h light/dark cycle. All procedures were approved by the Animal Care and Use Committee of the Technion – Israel Institute of Technology.

#### 4.1.2 Neuropixels probes and encapsulation

We used Neuropixels 1.0 (single-shank) and Neuropixels 2.0 (multi-shank) probes (IMEC)^62,63^. Because the external probe PCB and headstage interface are exposed during chronic recordings, we used a protective encapsulation to enable stable long-term implantation.

##### Chronic probe encapsulation

We designed a lightweight, low-cost 3D-printed encapsulation to protect Neuropixels probes during chronic implantation (Supplementary Fig. **??**). The encapsulation includes an insertion tool to facilitate stable probe positioning during surgery and, when needed, to enable safe post-mortem extraction of the shanks for histological analysis with minimal tissue disruption. The design also supports probe reuse and accommodates the headstage electronics for rapid connection/disconnection of the recording cable.

##### Probe trajectory planning

Probe insertion trajectories were planned preoperatively using the Neuropixels Trajectory Explorer toolbox^64^, which overlays candidate shank paths on the Allen Mouse Common Coordinate Framework^65^. This allowed us to select insertion angles that maximized coverage of the SUBC and MEC while avoiding ventricles and minimizing traversal of overlying cortex. Planned trajectories were then translated into stereotaxic coordinates and used to set the dorsoventral, anteroposterior and mediolateral angles on the micromanipulator during surgery.

#### 4.1.3 Surgical procedures

Mice were chronically implanted with a Neuropixels 1.0 or 2.0 probe targeting the SUBC and MEC. Before surgery, probes were mounted in the 3D-printed encapsulation and a thin silver wire was soldered to ground. Anesthesia was induced with isoflurane (4%) and maintained at 1.5–2% throughout surgery. Mice were secured in a stereotaxic frame (Kopf Instruments), and body temperature was maintained at 37 °C using closed-loop heating.

Preoperative treatments included analgesia (0.05 mg/kg buprenorphine), eye protection (DuraTears), and local anesthesia (lidocaine, 3 mg/kg, s.c.) before incision. The scalp was shaved and disinfected, and a 1-mm craniotomy was performed over the right hemisphere. After dura removal, probes were lowered stereotaxically using the coordinates and angles listed in Table **??**, traversing cortex (VI or TEA) to target SUBC and MEC. The probe ground wire was connected to a ground wire from a skull screw implanted in the left hemisphere. The encapsulation was secured using Metabond (Parkell, Edgewood, NY), and exposed wires and the incision site were covered with light-cured dental cement.

#### 4.1.4 Histological verification

Long-term Neuropixels recordings complicate post hoc verification of probe placement because dye coatings can fade and probe tracks can be difficult to resolve after extraction. To improve track visibility, we used paraffin embedding and, when feasible, left the probe shank *in situ* for sectioning with the tissue; sections were cut perpendicular to the shank when possible.

After completion of all recordings, mice were deeply anesthetized with isoflurane (5%) and transcardially perfused with 4% paraformaldehyde (PFA) in phosphate-buffered saline (PBS). Brains were extracted and post-fixed in 4% PFA at 4°C for 24 h. Depending on the preparation, the probe shank was removed after perfusion or left in place for subsequent sectioning. Samples were dehydrated and cleared using a tissue processor (Leica TP1020, Germany), embedded in Paraplast paraffin (Leica) using a Histocore Arcadia H embedding system (Leica), and sectioned at 5–10 *µ*m thickness at 50–100 *µ*m intervals using a rotary microtome (Leica RM2265, Germany). Sections were floated on a 36–38°C water bath, mounted onto adhesive glass slides (Leica X-tra), and dried overnight at 37°C.

Sections were deparaffinized in xylene, rehydrated through a graded ethanol series, and stained with hematoxylin and eosin (H&E) or Nissl stain (Table **??**). Slides were digitized and anatomical regions were identified by comparison to the Paxinos and Franklin mouse brain stereotaxic atlas^66^. Probe tracks and electrode landing sites were reconstructed by matching digitized sections to atlas plates. Representative histological sections from animals 1120, 1170, 7394 and 1150 are shown in Supplementary Fig. **??**b with atlas^66^ overlays.

### 4.2 Experimental and Behavioral Setup

#### 4.2.1 Experimental setup and tracking

To enable unrestricted movement in complex-shaped arenas, a motorized XY stage was suspended above a 120 ×120 cm experimental arena to keep the tethered recording cable directly above the animal and prevent entanglement (Supplementary Fig. **??**d). Animal rotations were accommodated by thin twisted tether wires, which were manually untwisted after each trial. The stage carried a camera (15 Hz; 66 ms per frame) and a closed-loop controller that used DeepLabCut^67^ to track a defined body point (tail–body junction) and dynamically reposition the stage to keep this point centered in the camera field of view.

Absolute position was obtained by composing the stage XY coordinates with the within–field-of-view tracked position, yielding high-resolution behavioral tracking. The same stage-mounted view enabled head-direction estimation using DeepLabCut-based landmark detection (the ears and nose). When needed, the XY stage could be manually controlled via a MATLAB interface to gently bias animals toward undersampled regions of the arena. A secondary fixed overhead camera (8.3 Hz; 120 ms per frame) provided independent monitoring for experimental oversight and validation.

Synchronization between electrophysiology recordings and video streams was achieved with external sync pulses delivered every 15 ± 2 s, logged both as camera frame markers and as digital events alongside the analog neural signals.

### 4.3 Neural Recordings and Preprocessing

#### 4.3.1 Neural Recordings

Neural signals were recorded continuously at 30 kHz using a National Instruments system (NI PXIe-6374). For Neuropixels 1.0, local field potentials (LFPs) and spikes were processed separately; for Neuropixels 2.0, LFPs were computed from spike data via digital filtering. Spike sorting was performed with Kilosort 2.5^63^, followed by automated curation with Bombcell^68^ to classify clusters and exclude noise and unstable units.

#### 4.3.2 Preprocessing

Behavioral variables (position and head direction) were extracted from DeepLabCut outputs^67^ and resampled/aligned to the neural time base using the recorded synchronization events. Sessions were included only if they passed quality-control criteria, including minimum unit yield and sufficient behavioral coverage of the arena.

### 4.4 Single Cell Analysis

#### 4.4.1 Rate maps

Spatial firing-rate maps (rate maps) were computed by discretizing the arena into 4 × 4 cm spatial bins and assigning each detected spike to the corresponding bin. Occupancy was estimated as the number of video frames in which the animal’s head position fell within a bin, multiplied by the frame period (40 ms), and firing rate was computed as spike count divided by occupancy time. Bins with *<* 0.2 s occupancy or *<* 2 spikes were excluded.

Units classified by Kilosort as multi-unit activity (MUA), flagged by Bombcell^68^ as noise, with *<* 500 spikes in the session, or with mean firing rates *<* 0.1 Hz or *>* 50 Hz were excluded from further analyses. Time points were excluded if the animal’s speed was *<* 2 cm/s (rest) or *>* 20 cm/s (implausible under tethering), or if DeepLabCut detection likelihood was *<* 50%. Rate maps were smoothed with a 2D Gaussian kernel (*σ* = 2 bins).

#### 4.4.2 Cell classification

##### Spatial information

We calculated spatial information (SI; bits per spike) for each cell as

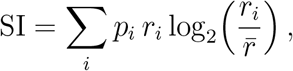

where *r*_*i*_ is the firing rate in spatial bin *i, p*_*i*_ is the occupancy probability of bin *i*, and 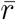 is the mean firing rate across bins. Statistical significance was assessed by comparing the observed SI against a null distribution generated via 1,000 random circular time shifts of the spike train within the corresponding 20-min room segment (excluding a 10-s margin at the start and end). The SI was considered significant if it exceeded the 95th percentile of this null distribution (*P <* 0.05).

##### Head direction

For each neuron, we computed a head-direction tuning curve by discretizing head direction into 6° bins. For bin *j*, firing rate was

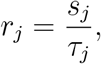

where *s*_*j*_ is the number of spikes fired when head direction fell within bin *j*, and *τ*_*j*_ is the total time spent in bin *j*. The peak firing rate was *r*_peak_ = max_*j*_(*r*_*j*_).

Directionality was quantified using the mean resultant vector length (Rayleigh vector length) across all angular bins,

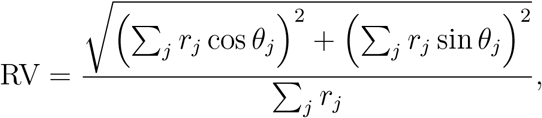

where *θ*_*j*_ is the center angle of bin *j*. The preferred head direction was

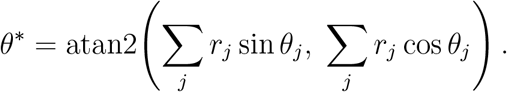

Statistical significance was assessed by comparing the observed Rayleigh score RV against a null distribution generated via 1,000 random circular time shifts of the spike train relative to behavior. Neurons were classified as HD cells if they met both criteria: (i) RV ≥ 0.3 and (ii) *P <* 0.05 relative to the null distribution^16,69^.

##### Allocentric border-vector cells

For each cell and behavioral epoch, we computed at each time step (i) the distance from the animal’s head to the nearest arena border *d*, and (ii) the allocentric angle from the head to the nearest wall *θ*. Angles were discretized into 10° bins. Distances were discretized into non-uniform bins (0–2 cm, 2–5 cm, 5–15 cm, 15–25 cm, and 25–40 cm) to provide higher resolution near walls.

For each (distance bin *k*, angle bin *m*) pair, firing rate was

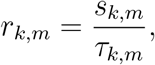

where *s*_*k,m*_ is the number of spikes fired while the animal occupied bin (*k, m*), and *τ*_*k,m*_ is the total time spent in that bin. Bins with *τ*_*k,m*_ *<* 0.2 s were set to zero (*r*_*k,m*_ = 0), and the map was smoothed with a Gaussian kernel (*σ* = 2 bins). The preferred wall distance and angle were defined by

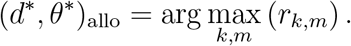

Statistical significance was assessed by comparing the observed Rayleigh score against a null distribution generated via 1,000 random circular shifts of the spike times (recomputing the allocentric angle tuning score for each shuffle). Cells were classified as allocentric border-vector cells if the score exceeded the 95th percentile of this null distribution (*P <* 0.05).

##### Egocentric border-vector cells

For each cell and each video frame, we computed the ego-centric distance to the nearest wall as a function of wall bearing relative to the animal’s head direction. Egocentric angles *θ*_ego_ were discretized into 10° bins spanning −180° to 180°. For each time point, distances were obtained by ray-casting from the head position at angles *θ*_ego_ and taking the closest wall intersection, then discretized into non-uniform bins (0–2 cm, 2–5 cm, 5–15 cm, 15–25 cm, and 25–40 cm).

For each (distance bin *k*, angle bin *m*) pair, firing rate was

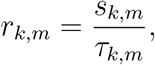

where *s*_*k,m*_ is the number of spikes fired while the animal occupied bin (*k, m*), and *τ*_*k,m*_ is the total time spent in that bin. Bins with *τ*_*k,m*_ *<* 0.2 s were set to zero and the map was smoothed with a Gaussian kernel (*σ* = 2 bins). The preferred egocentric distance and angle were

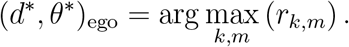

Cells were classified as EBCs if their tuning was significant relative to a null distribution generated via 1,000 random circular time shifts (*P* ≤ 0.05), identical to the allocentric border cell analysis.

##### Speed cell

Speed modulation was assessed by relating firing rate to running speed estimated from tracking. Instantaneous speed (cm/s) was computed in 1-s non-overlapping bins from the position trace; missing values were linearly interpolated, and negative values were set to zero. Speed bins (1 cm/s width) were defined across the observed range. For each speed bin, the mean firing rate was computed as the mean spike rate (spikes/s) across all 1-s time bins whose speeds fell within the interval.

The relationship between mean firing rate and speed was quantified using ordinary least-squares linear regression, and *R*^2^ was used as an index of speed modulation. Statistical significance was assessed by comparing the observed *R*^2^ against a null distribution generated via 1,000 random circular shifts of the spike times relative to behavior, recomputing *R*^2^ for each shuffle to generate a null distribution. Neurons were classified as speed-modulated if their observed *R*^2^ exceeded the 95th percentile of this null distribution (*P* ≤ 0.05) and met an effect-size criterion of *R*^2^ ≥ 0.6. Neurons that met these criteria and exhibited a positive regression slope were classified as putative speed cells.

#### 4.4.3 Cross-room maintenance of functional cell classes

To quantify whether neurons retained their functional classification across environments, we analyzed sessions consisting of three epochs (room A, room B, and A^′^ when available) and compared class membership between A vs. B and A vs. A′. Cells were classified within each epoch using the metrics described above (SI; Rayleigh vector length for allocentric border, ego-centric border, and HD tuning; and linear speed modulation quantified by *R*^2^), with significance determined against shuffle-based null distributions.

For each session and functional class, we counted the number of cells significant in epoch A (*n*_sig,A_) and computed how many of these were also significant in epoch B (*n*_maint,AB_) and in epoch A′ (*n*_maint,AA_*′*). Maintenance fractions were

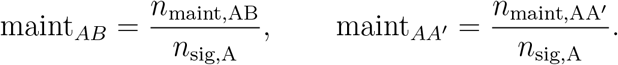

Thus, “retained classification” indicates that a neuron met the class criteria (metric threshold and shuffle-based significance) in both epochs being compared, conditional on being significant in epoch A. Directional classes used a Rayleigh vector-length threshold RV ≥ 0.3, and speed modulation additionally required *R*^2^ ≥ 0.6, in both cases in addition to shuffle-based significance.

#### 4.4.4 Cross-room changes in preferred direction among maintained cells

For directional cell classes that retained their functional identity across rooms (ABCs, EBCs, HD cells; see above), we quantified changes in preferred orientation between epochs A–B and A–A′. For each neuron and epoch, we first computed its preferred allocentric border, egocentric border, or head-direction angle from the corresponding tuning curve as the circular mean direction of the Rayleigh vector used for classification. For every neuron that was significantly directional in both epochs being compared (i.e. met the Rayleigh vector-length threshold RV ≥ 0.3 and passed the shuffle-based significance test in both A and the second epoch), we calculated the absolute difference between its preferred angles across rooms (epoch B or A′ minus epoch A).

Angle differences were aggregated across sessions for each directional class and comparison (A–B, A–A′) and summarized as polar histograms using fixed-width bins (bin centers at 0, 30, …, 180°) (Supplementary Fig. **??**f).

### 4.5 Position Decoding Model

We developed a decoding model based on population dynamics to obtain a quantitative description of mapping across rooms (Fig. 3a). We chose a model class that is highly expressive and recurrent so that it could, in principle, capture arbitrary history-dependent nonlinear relationships between firing patterns and position within each room. This allowed us to test whether the between-rooms relationship remained simple even when the within-room decoding model was maximally general. Neural and behavioral time series were aligned to a common 20 Hz sampling grid, with population activity computed as the raw spike count within a 250 ms sliding window centered at each time step.

#### Notations

Let *t* ∈ {1, …, *T*} be the relative time step within a single sequence, where *T* denotes sequence length. Let **R** ∈ ℝ^*N* ×*T*^ denote a sequence of population activity, where *N* is the number of cells, and the *t*-th column is a population vector **r**_*t*_ ∈ ℝ^*N*^. We take **P** ∈ ℝ^2×*T*^ to be the corresponding position trajectory, where **p**_*t*_ = (*x*_*t*_, *y*_*t*_)^⊤^,the *t*-th column, contains the *x* and *y* coordinates. The coordinate origin was defined at the top-left corner of the arena.

#### Model

We trained a neural-network decoder to predict position from population activity. The model factorizes into: (i) a shared temporal encoder *f* that maps neural activity to a low-dimensional latent trajectory, and (ii) room-specific decoder heads *g*_*c*_ (for room *c* ∈ {*A, B, A*^′^}) that map the shared latent trajectory to positions in the coordinate frame of each room (Fig. 3a). For a given sequence **R** sampled from room *c*:

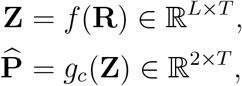

where *L* is the latent dimensionality. The temporal encoder, *f*, is a unidirectional recurrent long short-term memory (LSTM) layer^70^ (512 hidden units) followed by a multi-layer perceptron (MLP) network (two hidden layers of 256 and 128 units, ReLU (rectified linear unit) activation), yielding a latent trajectory **Z** = [**z**_1_, …, **z**_*T*_] ∈ ℝ^*L*×*T*^ where *L* = 64. For each room *c*, we trained a dedicated model to decode position from the shared latent coordinates per time step:

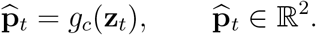

Each decoder head consisted of four hidden layers of 32, 32, 16, and 8 units, with ReLU activations and a final linear layer to produce the estimated position, 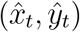.

#### Training

Sequence samples were constructed from *T* = 100 consecutive timestamps (spanning 5 s) with a stride of one timestamp (Δ*t* = 50 ms). Position coordinates were normalized to the [0, 1]^2^ interval via min-max scaling relative to room boundaries.

We trained the model to minimize the mean squared error (MSE) between predicted and observed positions. To enforce robust temporal integration while heavily penalizing errors at later timesteps that possess maximal sequence context, we computed an exponentially weighted loss over all time bins *t*, where 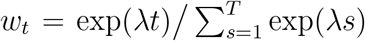 with *λ* = 0.01. The sequence-weighted prediction loss was

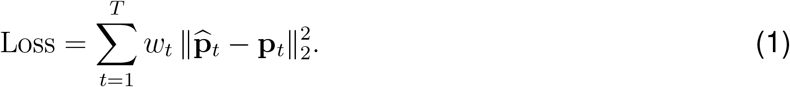

We computed the loss over the full sequence 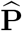, whereas we evaluated performance using only the final prediction 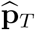. Training was performed in two stages to prevent the shared representation for the two visits to room A from biasing toward the repeated environment. In the first phase, the temporal encoder and room-specific heads for rooms *A* and *B* were trained jointly, excluding room *A*^′^. In the second phase, the shared modules were frozen, and only the decoder head for room *A*^′^ was trained. Model weights were optimized using Adam with a learning rate of 1 × 10^−3^, *p* = 0.1 dropout, and weight decay of 1 × 10^−4^. Early stopping was triggered if validation performance did not improve for 50 consecutive epochs (maximum: 500 epochs).

#### Cross-Validation

We evaluated model performance using 10-fold cross-validation with 10 repetitions. For each room, we partitioned the data into 10 contiguous temporal segments. In each fold, one segment served as the test set, while the final one-ninth of each remaining segment was reserved for validation (Supplementary Fig. **??**c). The remaining data constituted the training set. We prevented data leakage across overlapping temporal windows by using only sequences fully contained within a single contiguous segment. To make the results robust to the relative temporal alignment between rooms, we repeated the entire 10-fold procedure 10 times, each time circularly shifting the fold assignments of room B relative to room A (Supplementary Fig. **??**c). Overall, this yielded 10 folds × 10 repetitions = 100 trained models per session.

#### Evaluation

We evaluated test performance using only the final timestamp prediction 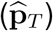 of each sample sequence, concatenating these predictions to reconstruct a continuous, full-length trajectory for each of the 10 cross-validation repetitions. We quantified decoding accuracy using two metrics, both computed as a pooled *R*^2^ across *x* and *y* coordinates: (i) **Mean** *R*^2^: The *R*^2^ was calculated for each repetition individually, then averaged across the repetitions. (ii) **Ensemble** *R*^2^: We first computed an “ensemble trajectory” (mean 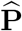) by averaging the predicted coordinates across the repetitions at each time step, then calculated the *R*^2^ of this mean trajectory against the true trajectory **P**. Averaging across repetitions improved the median decoder performance 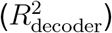 by 0.10 relative to individual models (Supplementary Fig. **??**f). Unless otherwise stated, reported performance metrics refer to the ensemble *R*^2^ and the corresponding Root Mean Squared Error (RMSE, in cm). Furthermore, all downstream geometric transformation analyses used the ensemble predictions to maximize stability.

#### 4.5.1 Latent space dimensionality and separability

To characterize the structure of the learned representations, we extracted latent trajectories (**Z**) by feeding the full dataset of population activity sequences through the shared temporal encoder, *f*. We quantified the dimensionality of the population code using Principal Component Analysis (PCA) on the concatenated latent vectors (**z**), determining the number of principal components required to explain 95% of the total variance (mean ± s.d. across models). For comparison, we applied identical PCA to the raw spike count vectors (**r**, model input).

To quantify geometric overlap between room representations, we assessed the linear separability of room identity. For each room pair, we trained a linear Support Vector Machine (SVM) to predict room identity from the first three principal components. To test for separability rather than generalization, we evaluated classification accuracy on the full dataset (all samples). We reported the misclassification rate (1 − accuracy) as a measure of overlap. Chance level was defined as the majority-class baseline (min(*n*_1_, *n*_2_)*/*(*n*_1_ + *n*_2_) ≈ 0.5). This analysis was repeated using the first three principal components of the raw neural activity.

#### 4.5.2 Comparison with Baseline Models

We compared our full architecture against two instantaneous (*T* = 1; Supplementary Fig. **??**h) baselines: (i) a **linear regression baseline** mapping instantaneous population activity (**r**) to spatial coordinates (**p**); and (ii) a **multilayer perceptron baseline** with three hidden layers of 512, 256, and 32 units, and ReLU activation. These models test the extent to which position can be decoded from instantaneous activity using simple linear or complex nonlinear transformations, serving as controls for the full architecture’s temporal integration.

### 4.6 Geometric Transformation Analysis

We quantified the geometric structure of spatial remapping by computing position correspondences as described below between room coordinate frames. We evaluated these transformations for all ordered room pairs (*A* → *A*^′^, *A* → *B, B* → *A*^′^).

#### 4.6.1 Deriving spatial correspondences

To ensure robust estimates, we established cross-environment correspondences using two complementary approaches: a continuous coordinate mapping derived from the neural network (NN) decoder (Section 4.5) and a discrete spatial bin assignment based on population-vector (PV) correlation (Section 4.6.1).

##### NN-based mapping

We utilized our trained model to infer the correspondence between spatial representations across rooms. For a given source room, we provided the decoder with neural activity recorded while the animal occupied that room, and first decoded the position in that room. We next presented the same activity pattern to the decoder while specifying the identity of a second, target room, thereby obtaining the position in the target room that was most consistent with that activity pattern under the learned model. This yielded a set of matched coordinate pairs 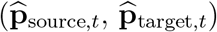, for each time step *t* (Fig. 3e, schematic; Supplementary Fig. **??**i).

##### PV-based mapping

For each spatial bin in each room (34 × 34 grid, occupancy ≥ 0.2 s), we computed the average firing rate of the population within that bin. We assigned each source-room spatial bin to the target-room bin that maximized the Pearson correlation between the population vectors of the two bins. Bins were considered matched if their maximum correlation exceeded 0.2. The strength of this mapping was evaluated by averaging PV correlations across all matched bin pairs.

#### 4.6.2 Significance testing of PV mapping

To determine whether the discrete spatial mappings exhibited non-random structure, we tested the statistical significance of the PV correlation against null distributions generated via 1,000 random data perturbations. Each perturbation type was designed to selectively disrupt specific relationships between rooms.

##### Cell-index shuffling

To disrupt neuron-identity correspondence across rooms, we randomly permuted cell identities between the two rooms before recomputing mapping metrics.

##### Spatial bin shuffling

To disrupt spatial structure while preserving other aspects of the data, we randomized the spatial arrangement of bins within the source room using (i) independent row/column shuffles, (ii) systematic shifts, or (iii) reorganization within defined spatial rings, and then recomputed the PV correlation. Across shuffle variants, the null hypothesis was that apparent mapping structure is expected by chance rather than reflecting a true transformation between rooms.

#### 4.6.3 Comparison of mapped coordinates

To directly compare the spatial transformations inferred by the NN decoder and PV analysis (Fig. 3e,f), we discretized the continuous NN decoder output to match the 34 × 34 bin resolution of the PV analysis. For each source-room spatial bin, we identified all decoded positions falling within that bin, computed the centroid of their corresponding target-room coordinates, and assigned it to the nearest target spatial bin.

##### Smoothness

To quantify the local continuity of the mapping (Fig. 3g), we used the graph Laplacian quadratic form, also known as the graph Dirichlet energy. We built a graph based on the spatial bins of the source room and then measured the Dirichlet energy of the target coordinates on this graph, relative to the source coordinates. By construction, this measure is small if the mapping preserves the same local neighbors. Formally, we constructed a graph with *V* nodes representing the spatial bins in the source room, and each node is connected to its 8 adjacent spatial neighboring bins. Denote **A** as the adjacency matrix indicating which nodes are connected (edge value = 1) and which are not (=0), and **D** as the degree matrix, where **D**_*i,i*_ = ∑_*j*_ *A*_*i,j*_. Following standard spectral graph theory^71^, we computed the normalized graph Laplacian 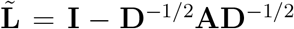. The coordinate column vectors **x, y** ∈ ℝ^*V*^ are treated as graph signals on this graph^72^. We defined the energy changes for the horizontal and vertical coordinates as:

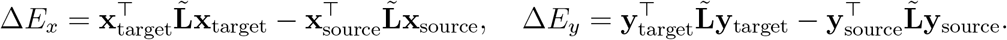

We combined these terms into a normalized smoothness score:

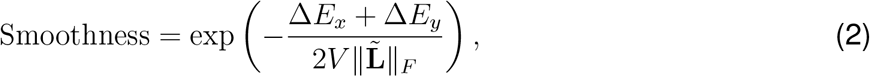

where 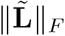 is the Frobenius norm of the normalized Laplacian matrix. We used the denominator 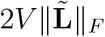 as a per-session normalization term to place energy changes on a comparable scale across sessions with varying numbers of bins and graph structures. The exponential transform was used to obtain a bounded, interpretable score. Higher values indicate that the mapping introduces less additional local variation relative to the source coordinate field. This metric was calculated for both the NN and PV mappings. Only sessions with at least 100 valid bins were included. Statistical significance was assessed by comparing the observed smoothness score against a null distribution generated via 1,000 random bin-to-bin mappings. Scores were considered significant if they exceeded the 95th percentile of this null distribution (one-sided test, *P <* 0.05).

#### 4.6.4 Affine Transformation estimation

We estimated the optimal global affine transformation mapping from the source room to the target room. We used least squares to fit an affine transformation to the position correspondences. We then decomposed the resulting mapping into interpretable components (rotation, reflection, stretching, and translation). The goodness of fit of the estimated transformation was quantified by the mean squared error (MSE) between the affine-transformed source coordinates and the corresponding target coordinates, as well as by the coefficient of determination 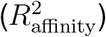. We hereafter refer to this 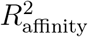 as the **affinity score**. High affinity scores indicate that the relationship between room representations is well-approximated by a global affine model.

##### Affine transformation decomposition

We decomposed the estimated affine transformations into explicit geometric components. In homogeneous coordinates, the transformation maps a point 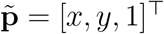 to 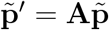, where **A** ∈ ℝ^3×3^ is defined as:

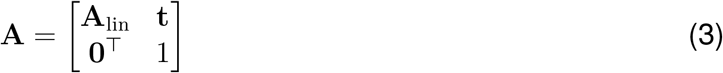

**A**_lin_ ∈ ℝ^2×2^ encodes linear transformations (rotation, reflection, scaling) and **t** ∈ ℝ^2^ represents translation.

##### Polar decomposition

We factored the linear component **A**_lin_ (Eq. 3) using the left polar decomposition:

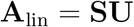

where **U** is an orthonormal matrix and **S** is a symmetric positive-definite scaling matrix. The left-polar form implies that scaling occurs along axes defined in the target frame. The determinant of **U** determined whether the transformation involved a rotation (det(**U**) *>* 0) or a reflection (det(**U**) *<* 0).

##### Rotation and reflection angles

For rotations (det(**U**) *>* 0), we computed the rotation angle:

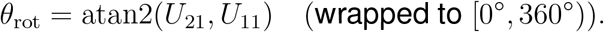

For reflections (det(**U**) *<* 0), we identified the reflection axis **v** = (*v*_*x*_, *v*_*y*_)^⊤^ as the eigenvector of **U** with eigenvalue +1. The axis orientation was computed as:

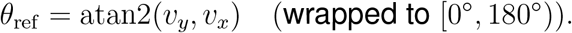

##### Anisotropic scaling

We decomposed the symmetric matrix **S** to obtain scaling factors *s*_1_ ≥ *s*_2_ *>* 0 and their corresponding orthogonal principal axes. These factors quantify expansion or compression along specific directions in the target frame.

#### 4.6.5 Significance of affinity scores

We compared the obtained affinity scores, 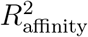, against a null distribution generated via 1,000 cyclic time-shifts of the target data relative to the source. This preserves temporal structure within each room while disrupting cross-room correspondence. Observed affinity scores were considered statistically significant if they exceeded the 95th percentile of the null distribution (*P <* 0.05). Additionally, we applied a minimum threshold of 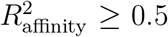 to classify a transformation as structurally affine.

### 4.7 Comparison of NN model and PV analysis

We compared the transformation parameters obtained from PV analysis (Section 4.6.1) to those obtained from the NN model. We included sessions meeting decoder and affinity criteria 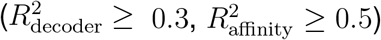 where both methods agreed on the transformation type (rotation or reflection) and yielded valid angle estimates (Fig. 4d).

We extracted rotation angles (*θ*_rot_), reflection angles (*θ*_ref_), and scaling factors (*s*_1_, *s*_2_) from the affine models fitted by both methods. Orientation parameters were standardized to [0°, 360°) for rotations and [0°, 180°) for reflection. We assessed consistency using circular-linear regression (evaluating slope and circular *R*^2^; see Circular regression analysis below) and quantified angular error using RMSE.

### 4.8 Transformation consistency across neural subpopulations

We retrained our model per subpopulation where neurons were partitioned by region (MEC, SUBC) or by functional class (ABC, EBC, HD, SI, Speed) as defined in single-cell analysis (Section 4.4). Cells with mixed selectivity contributed to multiple functional subgroups; subgroups with fewer than 10 cells were discarded. A majority of sessions achieved 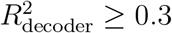. Angular consistency of the resulting transformations between subpopulations and the full ensemble was then evaluated (see Circular regression analysis below).

### 4.9 Circular regression analysis

We quantified the consistency of transformation parameters (rotation and reflection angles) across methods (PV vs. NN) and neural subpopulations, using circular regression framework as follows. We modeled the relationship as a directional fit *θ*_2_ ≈ *a* · *θ*_1_ + *b* by minimizing the circular loss

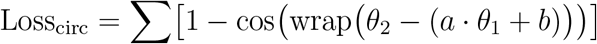

using the Nelder-Mead algorithm. For reflection axes (180° periodicity), angles were doubled prior to analysis, and the resulting intercept was halved. Given the nonlinear nature of this loss, statistical significance was assessed by comparing the circular *R*^2^ against a null distribution generated via 1,000 random permutations of *θ*_2_ across pairs.

### 4.10 Single-cell conformity analysis

To determine whether global geometric transformations emerge from coherent single-cell remapping, we analyzed directional tuning shifts in ABCs. We included rotation sessions that met the decoder and affinity criteria 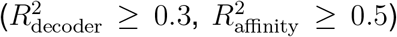 and passed the tuning-prominence filter (see below).

We computed the distribution of angular shifts for all ABCs (*n* ≥ 10 cells per session). Let *θ*^∗^ denote the preferred allocentric wall angle for an ABC (Section 4.4). The shift for cell *i* was

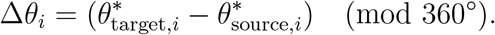

We assessed conformity and quantified the alignment between the session-level single-cell shift mode (*θ*_mode_) and the global affine rotation (*θ*_rot_). Define *θ*_mode_ as the circular peak of the smoothed Δ*θ*_*i*_ histogram (15° bins, Gaussian *σ* = 1 bin). Sessions were classified as conforming if the angular separation between *θ*_mode_ and *θ*_rot_ was ≤30°.

We evaluated conformity only in sessions exhibiting a clear population-level shift. Specifically, we required the peak prominence of the smoothed shift distribution to exceed its median baseline by a factor of ≥ 1.

### 4.11 Reflection axis alignment with room geometry

For transformations characterized as reflections (det(**U**) *<* 0), we evaluated the alignment of the reflection axis *θ*_ref_ relative to the shared wall orientation. We included sessions meeting decoder and affinity criteria 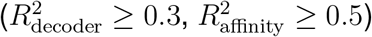.

### 4.12 General Methods

#### 4.12.1 Statistical analysis

We performed all analyses using Python 3.x with PyTorch^73^ for model training, NumPy and Pandas for data manipulation, and Weights & Biases (W&B)^74^ for experiment tracking. Matlab R2021a was used for for single-cell analyses, rate maps and PV analyses. We performed statistical comparisons across rooms and conditions using cross-validated estimates with appropriate corrections for multiple comparisons. We computed standard errors of the mean (SEM) across cross-validation folds to quantify uncertainty in performance estimates. All statistical tests were one-sided unless otherwise noted.

#### 4.12.2 Computational resources

Models were trained on NVIDIA GPUs. Typical training time for a single fold was approximately 5 ± 3 minutes depending on dataset size. All training runs were tracked using W&B, including hyperparameters, training curves, and performance metrics.

## Acknowledgements and declarations

**ACKNOWLEDGMENTS**

The work was funded by an NSF-BSF-NIH Computational Neuroscience (CRCNS) grant to D.D. and V.B. (NSF 2011509 BSF 2019807 NIMH 1 R01 MH125544-01), by Israel Science Foundation grants to H.B. (ISF 2418/24) and DD (ISF 2813/21), by a Rappaport Institute Collaborative research grant to D.D. and by the Prince Center for the Aging Brain to D.D. The authors thank all members of their labs for their support.

## AUTHOR CONTRIBUTIONS

Conceptualization, S.A., D.D., G.M., H.B. and D.Z.; Methodology, S.A., D.Z., D.D., H.B. and G.M.; Experimentation, S.A., G.T., S.L., O.R., and S.K.; Formal analysis, S.A. and D.Z.; Writing— original draft, S.A. and D.Z.; Writing—review & editing, H.B., D.D., G.M. and V.B.; Funding acquisition, D.D, V.B. and H.B.; Supervision, D.D., H.B. and G.M.

## DECLARATION OF GENERATIVE AI AND AI-ASSISTED TECHNOLOGIES

During the preparation of this work, the authors used Gemini, ChatGPT and Perplexity to assist with language improvement and used Perplexity and Claude to assist with limited code generation and code refinement. After using these tools or services, the authors reviewed and edited the content as needed and take full responsibility for the content of the publication.

## RESOURCE AVAILABILITY

### Lead contact

Requests for further information and resources should be directed to and will be fulfilled by Dori Derdikman and Hadas Benisty.

### Materials availability

This study did not generate new materials.

### Data and code availability

- Data is available upon request.
- Code for model training and evaluation is available at https://github.com/BenistyLab/population-decoder-remapping.
- Any additional information required to reanalyze the data reported in this paper is available from the lead contacts upon request.

