## Supplemental Materials for "Spatial remapping in the subicular complex and entorhinal cortex follows low-dimensional geometric principles" (from another version of this preprint: version 1, 2026-05-13)

### Supplementary Information and Tables for Abramson et al.

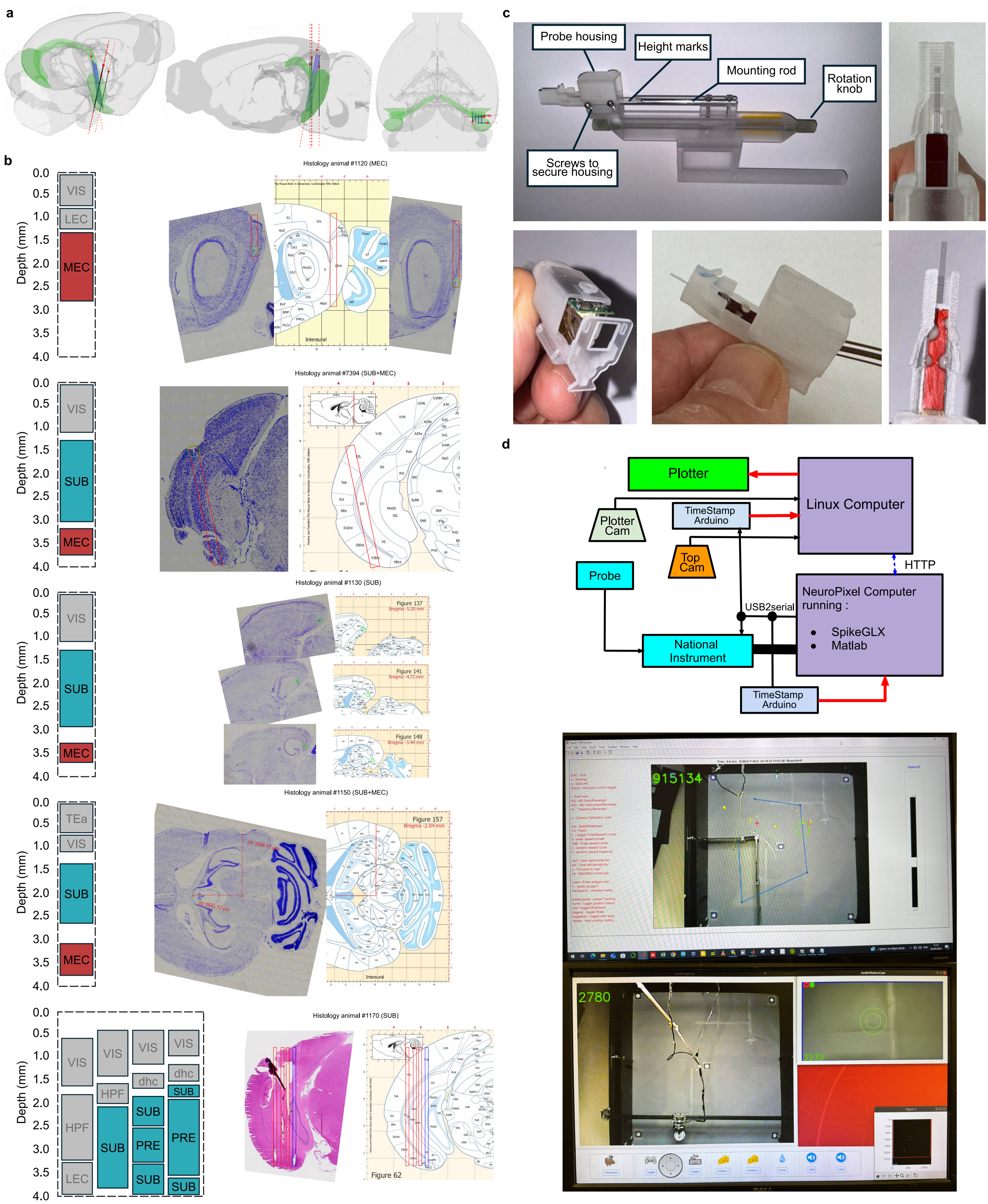

Figure S1: **Supplementary Figure S1.** Recording geometry, histology and setup. **(a)** Neuropixels probe trajectories shown (left to right) in isometric, coronal and horizontal atlas views, [?] .Black lines, Neuropixels 1.0 probes; blue lines, Neuropixels 2.0 probes. **(b)** Histology. Sagittal section from mouse 1120 (top left) showing two probe tracks (green circles) and the corresponding reconstructed probe locations in the atlas[?] (red outline). Coronal section from mouse 7394 (top middle) showing the insertion point and angle (green circle) and the reconstructed probe trajectory in the atlas (red outline). Horizontal section from mouse 1130 (top right) showing probe tracks (green circles) and their corresponding atlas locations. Coronal section from mouse 1170 (bottom left) with a Neuropixels 2.0 four-shank probe showing the reconstructed locations of the shanks (red and blue outlines) in the atlas. Horizontal section from mouse 1150 (bottom right) showing the insertion point and reconstructed probe location in the atlas. For each histology panel, the brain regions sampled by the probe are indicated to the right. **(c)** Custom Neuropixels encapsulation and insertion tool, shown with a Neuropixels 2.0 four-shank probe. **(d)** Experimental setup. Block diagram of the acquisition system (top); setup-control user interface showing the top-camera and overhead moving-camera views (middle); and Matlab user interface (bottom) showing the arena with a planned trajectory (blue) and the mouse's location and head direction (green circle and red arrow).

2

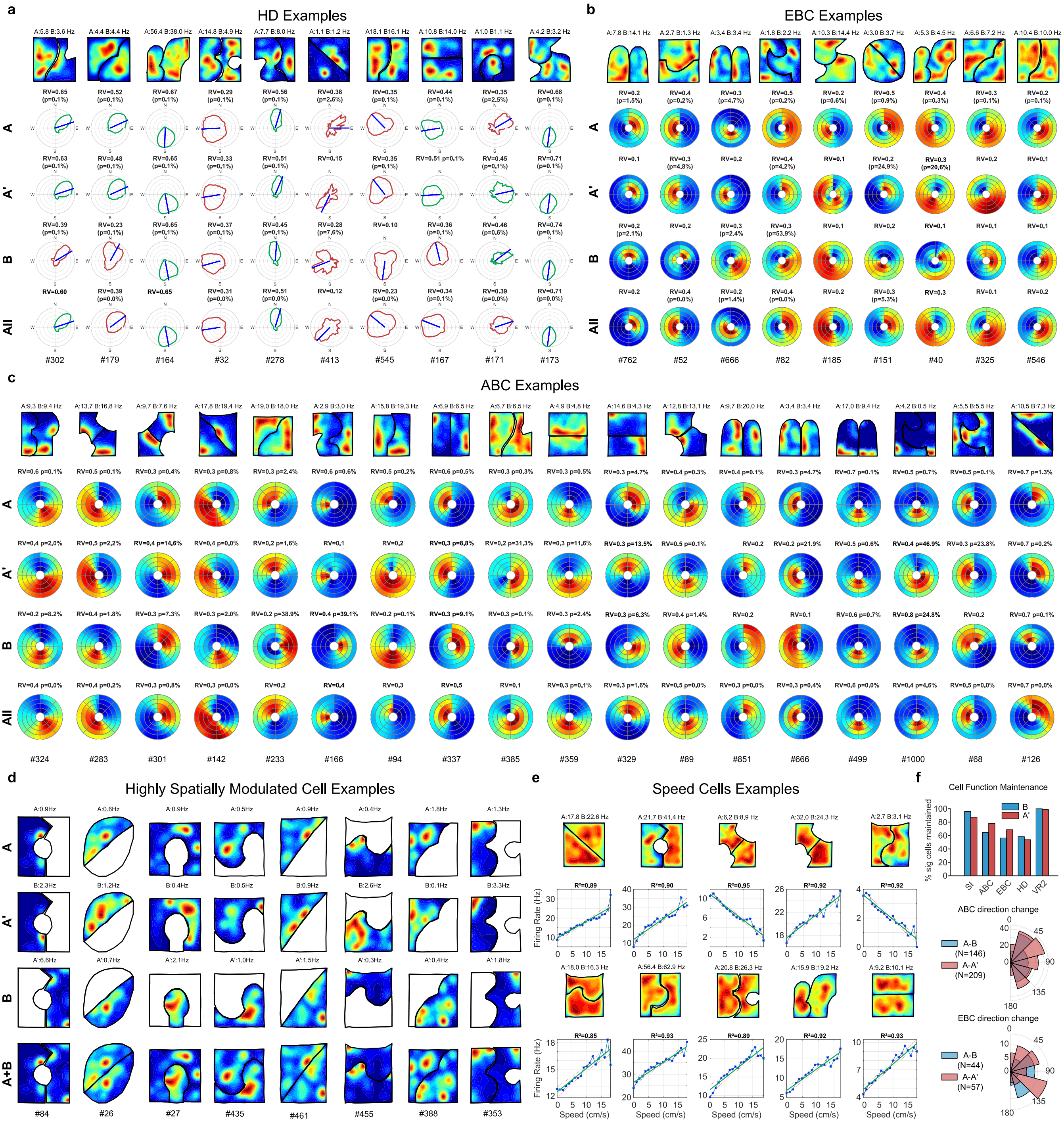

a
HD Examples
b
EBC Examples
c
ABC Examples
d
Highly Spatially Modulated Cell Examples
e
Speed Cells Examples
f
Cell Function Maintenance
% sig cells maintained
SI
ABC
EBC
HD
VR2
ABC direction change
A-B (N=146)
A-A' (N=209)
EBC direction change
A-B (N=44)
A-A' (N=57)
Firing Rate (Hz)
Speed (cm/s)

Figure S2: **Supplementary Figure S2.** Examples of functionally defined cell types and their stability across rooms. **(a)** Example head-direction (HD) cells. The top row shows spatial rate maps; rows 2–4 show directional tuning curves in rooms A, A′ and B; the bottom row shows tuning across the full session. Cell IDs are shown below each column. **(b)** Example egocentric border cells (EBCs). The top row shows spatial rate maps; rows 2–4 show egocentric polar maps in rooms A, A′ and B; the bottom row shows the egocentric map for the full session. Polar plots are shown in egocentric coordinates, with the top indicating the animal's forward direction and radius indicating distance from the wall (0–2, 2–5, 5–15 and 15–40 cm). Titles report Rayleigh vector magnitude and significance ($p$-value). Cell IDs are shown below each column. **(c)** Example allocentric border cells (ABCs). The top row shows allocentric rate maps; rows 2–4 show allocentric polar maps in rooms A, A′ and B; the bottom row shows the allocentric map for the full session. Polar plots are shown in allocentric coordinates, with the top indicating North and radius indicating distance from the wall (0–2, 2–5, 5–15 and 15–40 cm). Titles report Rayleigh vector magnitude and significance. Cell IDs are shown below each column. **(d)** Example highly spatially modulated cells. Rows 2–4 show spatial rate maps in rooms A, A′ and B; the bottom row shows the map for the full session. Titles report the maximum firing rate. Cell IDs are shown below each column. **(e)** Example speed cells. For each cell, the top panel shows the spatial rate map and the bottom panel shows firing rate as a function of running speed. The green line indicates the regression fit, and orange dashed lines indicate error bounds. **(f)** Functional classes and maintenance across rooms. The bar plot (top) shows the fraction of cells that preserved their functional category between rooms. Blue bars indicate maintenance between A and B, and red bars indicate maintenance between A and A′. Groups correspond to spatially informative (SI) cells, allocentric border cells (ABCs), egocentric border cells (EBCs), head-direction (HD) cells and speed cells. Polar plots (middle, ABC; bottom, EBC) show the absolute change in preferred Rayleigh angle for cells that maintained their function; blue indicates the A–B comparison and red indicates the A–A′ comparison.

3

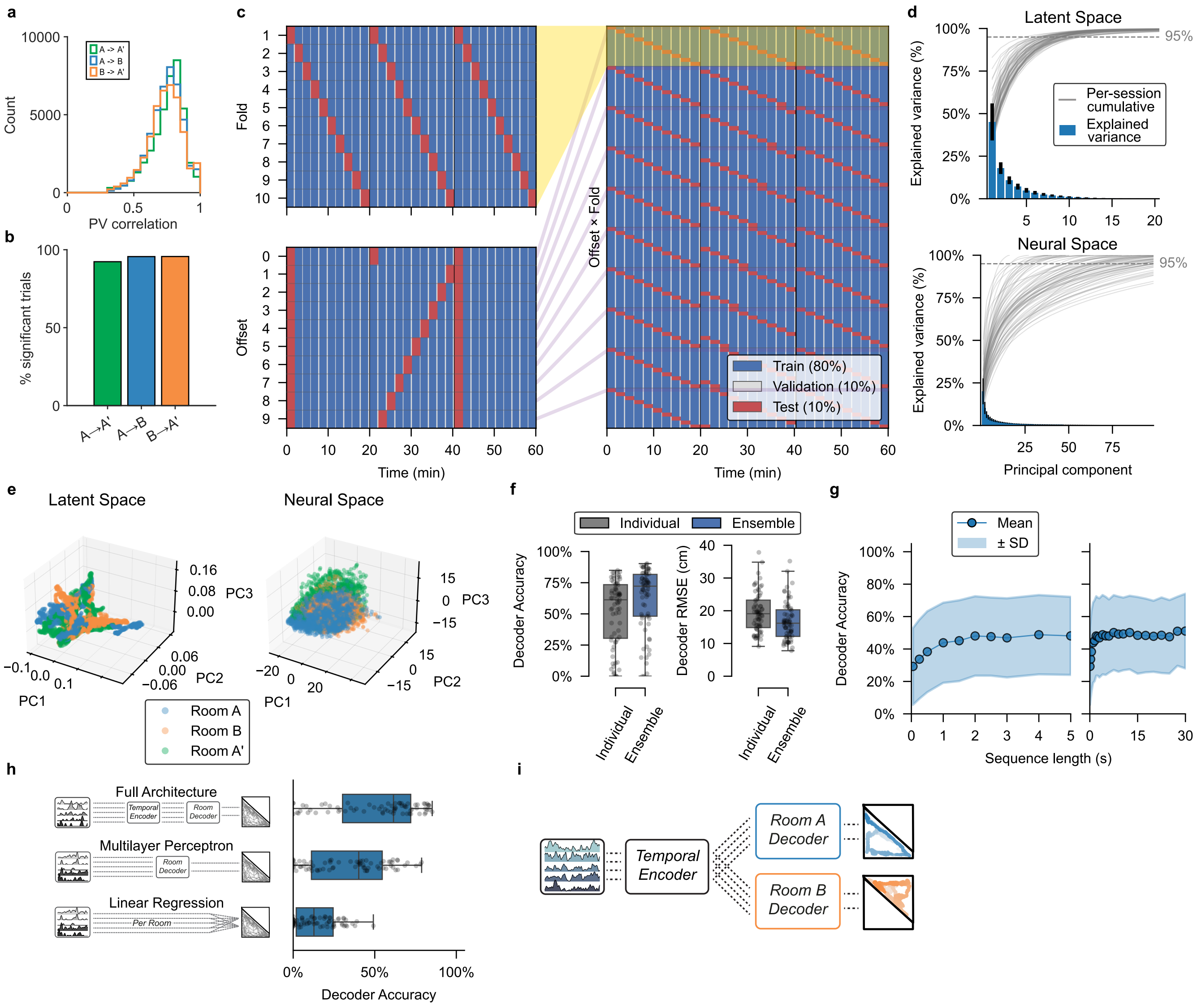
a
b
c
d
e
f
g
h
i
PV correlation
Count
% significant trials
Fold
Offset
Offset × Fold
Time (min)
Train (80%)
Validation (10%)
Test (10%)
Latent Space
Neural Space
Explained variance (%)
Per-session cumulative
Explained variance
95%
Principal component
PC1
PC2
PC3
Room A
Room B
Room A'
Individual
Ensemble
Decoder Accuracy
Decoder RMSE (cm)
Mean
± SD
Sequence length (s)
Full Architecture
Multilayer Perceptron
Linear Regression
Temporal Encoder
Room Decoder
Per Room
Room A Decoder
Room B Decoder

Figure S3: **Supplementary Figure S3.** Characterization of the latent space and validation of decoder performance. **(a)** Distribution of bins by their maximum PV correlation for room comparisons A–A′, A–B, and B–A′. **(b)** Percentage of trials with significant mean maximum PV correlation relative to shuffled-cell controls for room comparisons A–A′, A–B, and B–A′. **(c)** Cross-validation scheme. Schematic of the 10-fold cross-validation procedure. A continuous recording session is partitioned into temporally separated training (80%), validation (10%), and test (10%) segments, with rolling temporal offsets across repetitions to ensure evaluation is not driven by temporal autocorrelation. **(d)** Cumulative variance explained by principal components (PCs) for the latent space (top) versus raw neural activity (bottom). The latent representation requires substantially fewer components to reach 95% variance ($9 \pm 2$ PCs) compared to raw activity ($71 \pm 37$ PCs), indicating significant dimensionality reduction (mean $\pm$ s.d. across $N = 71$ sessions). **(e)** PCA visualization of room invariance. First three PCs of the latent space (left) and raw activity (right) for a representative single model, colored by room identity (blue: A; orange: B; green: $A'$). In the latent space and the raw neural activity, representations strongly overlap. Across all sessions, linear classification of room identity from these 3 PCs yields near-chance misclassification rates (latent: $0.39 \pm 0.01$; raw: $0.40 \pm 0.01$; mean $\pm$ s.d.; pairwise chance = 0.5), confirming that the latent code is effectively room-invariant. **(f)** Ensemble decoding stability. Boxplots comparing individual model repetitions (gray) with the ensemble average (blue). Ensembling improves median $R^2_{\text{decoder}}$ by $\sim$0.10 (individual: 0.61; ensemble: 0.72) and reduces median position error (RMSE) by $\sim$2.6 cm (individual: 19.1 cm; ensemble: 16.2 cm), demonstrating enhanced stability and accuracy ($N = 71$ sessions). **(g)** Decoder accuracy as a function of sequence length. Accuracy improves rapidly as temporal context increases up to a few seconds before plateauing, highlighting the importance of temporal integration (mean $\pm$ s.e.m. across $N = 71$ sessions). **(h)** Comparison to baseline models ($N = 71$ sessions). Boxplots demonstrate that the full architecture (median $R^2_{\text{decoder}} = 0.61$) significantly outperforms an instantaneous multilayer perceptron network (0.40) and linear regression (0.13), further supporting the necessity of temporal integration for decoding the population code. **(i)** Schematic of the cross-decoding framework. Neural population activity maps through a shared temporal encoder to a common latent state, which is then read out by room-specific decoders to infer cross-room correspondences.

4

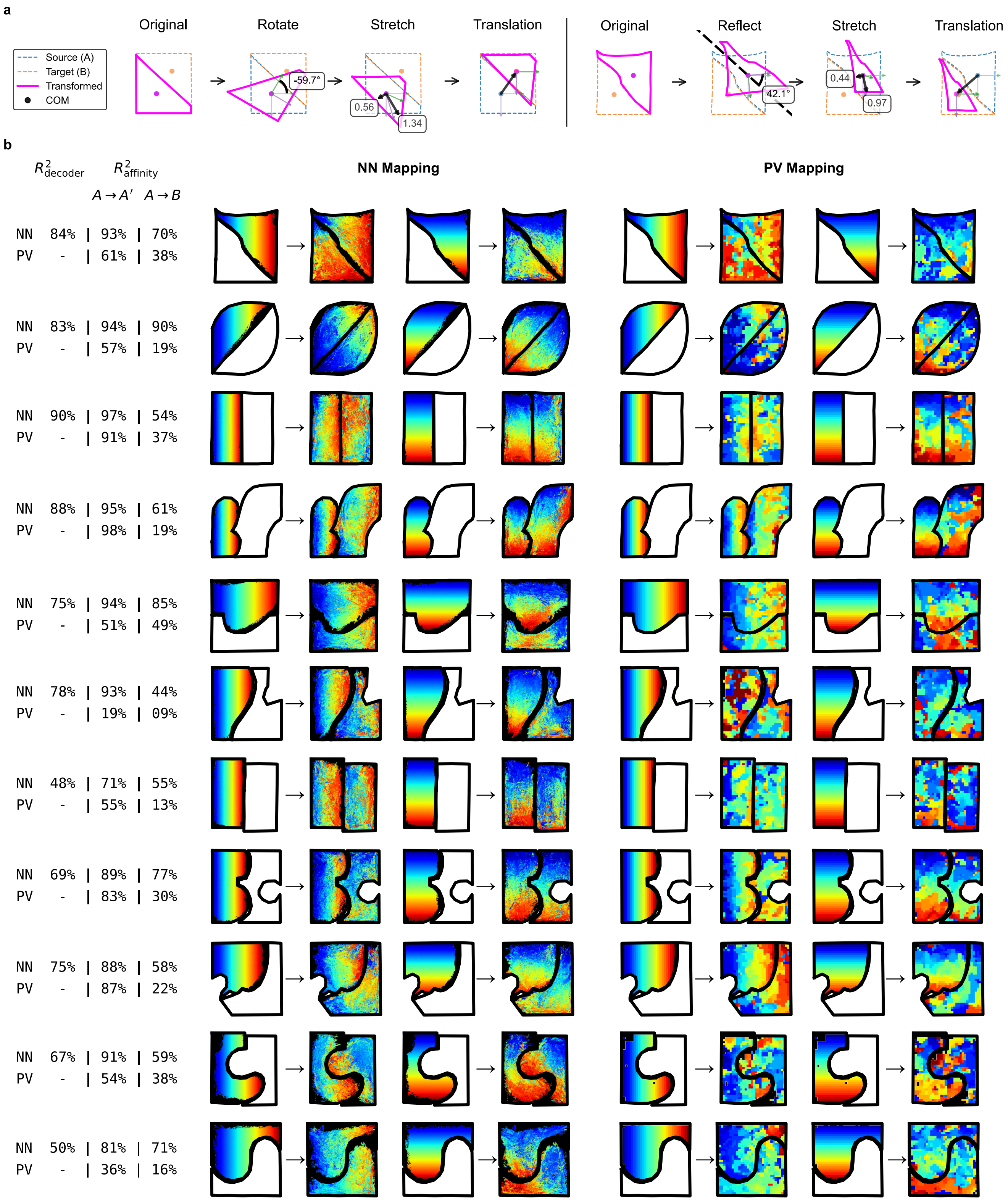
a
Original
Rotate
Stretch
Translation
Original
Reflect
Stretch
Translation
Source (A)
Target (B)
Transformed
COM
-59.7°
0.56
1.34
42.1°
0.44
0.97
b
$R^2_{decoder}$
$R^2_{affinity}$
$A \to A'$
$A \to B$
NN Mapping
PV Mapping
NN 84% | 93% | 70%
PV - | 61% | 38%
NN 83% | 94% | 90%
PV - | 57% | 19%
NN 90% | 97% | 54%
PV - | 91% | 37%
NN 88% | 95% | 61%
PV - | 98% | 19%
NN 75% | 94% | 85%
PV - | 51% | 49%
NN 78% | 93% | 44%
PV - | 19% | 09%
NN 48% | 71% | 55%
PV - | 55% | 13%
NN 69% | 89% | 77%
PV - | 83% | 30%
NN 75% | 88% | 58%
PV - | 87% | 22%
NN 67% | 91% | 59%
PV - | 54% | 38%
NN 50% | 81% | 71%
PV - | 36% | 16%

Figure S4: **Supplementary Figure S4.** Decomposition and global structure of affine spatial transformations. **(a)** Affine decomposition examples. Representative decompositions of cross-room mapping into elementary geometric operations. The source room (blue dashed contour; room A) is sequentially transformed (magenta contour) to align with the target room (orange dashed contour; room B), dots denote the center of mass. Left: a transformation dominated by rotation ($-59.7°$), anisotropic scaling (factors 0.56 and 1.34), and translation. Right: a transformation comprising reflection (dashed diagonal axis, $42.1°$), anisotropic scaling (factors 0.44 and 0.97), and translation. These examples demonstrate the decomposition of complex population-level remapping into a compact set of basic affine operations. **(b)** Gallery of cross-room mappings for representative sessions. Comparison of global mapping structure inferred by neural network decoders (NN Mapping, center) and population vectors (PV Mapping, right). Leftmost columns report session metrics: decoder accuracy ($R^2_{\text{decoder}}$) and affinity scores ($R^2_{\text{affinity}}$) for $A \to A'$ and $A \to B$. Within each method block, horizontal ($x$, left pair) and vertical ($y$, right pair) spatial gradients in the source room are projected onto the target room. NN mappings yield smooth, continuous gradient transformations (e.g., rotated, curved, or compressed bands), reflecting robust geometric correspondence across environments. PV mappings exhibit lower spatial precision, producing noisy coordinate assignments, yet preserve a similar global structure. Thus, while both methods capture related spatial transformations, NN decoding resolves the mapping with greater continuity and precision.

Table 1: **Recording region by animal.** Number of recording sessions per animal in SUBC, MEC, or sessions spanning both regions (SUBC+MEC).

| | **Region** | | |
|---|---|---|---|
| **Animal#** | **SUBC** | **MEC** | **SUBC+MEC** |
| 1120 | | 26 | |
| 1130 | 1 | | |
| 1140 | 3 | | |
| 1150 | | | 13 |
| 1170 | 24 | | |
| 7394 | | | 4 |

Table 2: **Probe implantation coordinates.** AP, anterior–posterior; ML, medio–lateral; DV, dorso–ventral.

| Animal | Neuropixels | AP (μm) | ML (μm) | Angle (deg) | DV (μm) |
|---|---|---|---|---|---|
| 1120 | 1.0 | −4500 | 4024 | 75H | 3500 |
| 1130 | 1.0 | −4000 | 3730 | 75H | 3500 |
| 1140 | 1.0 | −3900 | 4120 | 75H | 4200 |
| 1150 | 1.0 | −3900 | 4120 | 75H | 4200 |
| 7394 | 1.0 | −4300 | 4024 | 75H | 3500 |
| 1170 | 2.0 | −4350 | 2800 | 80H/10V / 200R | 4200 |

Table 3: **Staining protocol.** Protocol steps for hematoxylin and eosin (H&E) and Nissl staining.

| | **H&E** | **Nissl** |
|---|---|---|
| **Deparaffinization** | Xylene × 5 min ( ×3 ) | |
| **Re-hydration** | EtOH 100% × 1 min ( ×3 ) | |
| | EtOH 95% × 1 min | |
| | EtOH 80% × 1 min | |
| | Rinse in running tap water × 1 min | |
| **Stain** | Hematoxylin (Papanicolaou Harris hematoxylin) × 5 min<br>Rinse in running tap water × 3 min<br>Differentiator (0.3% alcoholic HCl) × 2 dips<br>Rinse in running tap water × 5 min<br>EtOH 95% × 1 min<br>Eosin alcoholic × 5 min | Cresyl violet (warm up to 50 ℃) × 20 min (put to heat 10 min before)<br>Quick rinse in running tap water × 2 dips<br>Differentiate in EtOH 95% × 10 sec |
| **Dehydration** | EtOH 95% × 10 sec | |
| | EtOH 100% × 1 min ( ×3 ) | |
| | Xylene × 1 min ( ×3 ); hold in the last one | |
